# Topological decoding of grid cell activity via path lifting to covering spaces

**DOI:** 10.1101/2025.10.17.683158

**Authors:** Yuxing Jared Yao, Iris H.R. Yoon

**Affiliations:** Department of Mathematics and Computer Science, Wesleyan University; Program in Neuroscience and Behavior, Wesleyan University; Department of Mathematics and Statistics, Swarthmore College

## Abstract

High-dimensional neural activity often resides in a low-dimensional subspace, referred to as neural manifolds. Grid cells in the medial entorhinal cortex provide a periodic spatial code that is organized near a toroidal manifold, independent of the spatial environment. Due to the periodic nature of this code, it is unclear how the brain utilizes the toroidal manifold to understand its state in a spatial environment. We introduce a novel framework that decodes spatial information from grid cell activity using topology. Our approach uses topological data analysis to extract toroidal coordinates from grid cell population activity and employs path-lifting to reconstruct trajectories in physical space. The reconstructed paths differ from the original by an affine transformation. We validated the method on both continuous attractor network simulations and experimental recordings of grid cells, demonstrating that local trajectories can be reliably reconstructed from a single grid cell module without external position information or training data. These results suggest that co-modular grid cells contain sufficient information for path integration and suggest a potential computational mechanism for spatial navigation.

## 1. Introduction

Activity of a population of neurons often resides in a low-dimensional subspace called a neural manifold [8, 20, 22, 23, 30, 33, 43, 46] whose structure reflects the information encoded by the neurons. For example, the activity of head direction cells [41] are organized near a circle [8, 33]. Grid cells in the medial entorhinal cortex (MEC) exhibit a periodic hexagonal firing pattern that tiles the environment at regular intervals [22] and are organized into modules whose cells share scale and orientation but differ by fixed spatial phase offsets [22, 27]. The periodicity of a single-module of grid cell activity implies that the population activity is topologically organized around a torus ^1^ [5, 19]. Such organization has been captured by continuous attractor network (CAN) models of grid cells and has been observed in large-scale recordings [20].

Because multiple locations in a spatial environment elicit a similar response among co-modular grid cells, spatial locations are not uniquely encoded in the toroidal neural manifold of grid cells. This insight raises a central question: how much spatial information can be decoded from the activity of a single module of grid cells? Prior efforts to decode position from neural activity often relied on place cell dynamics [2, 12, 15, 37] or on combining phase differences across multiple grid modules [26, 38]. Other methods perform cumulative vector integration [6] or train deep models to map activity to position [18, 24, 28, 40, 45]. Theoretical analysis indicates that a single grid module may carry sufficient information to update its internal representation of position [6, 16, 25, 39], known as path-integration, though explicit computational demonstrations have been limited, constrained to multiple trials of one-dimensional settings [44] or relying on access to firing rate maps and training of deep neural networks [31].

In this study, we present a novel method for decoding movement trajectories from the activity of a single module of grid cells. The method builds on the insight that the topology of a stimulus space can be recovered directly from neural activity [10] and integrates tools from topological data analysis and path lifting in topology: Persistent cohomology reveals the toroidal structure of grid cell activity, circular and toroidal coordinates parametrize grid cell activity on this torus, and path lifting reconstructs the movement path in Euclidean space. To the authors’ knowledge, this is the first work integrating path lifting in topology into computational and applied settings. The approach differs from existing decoding work in that it only uses data from a single module of grid cells and that it doesn’t involve any training process. We validate the algorithm in both CAN-simulated and experimental datasets by showing that the reconstructed movement paths differ from the original by an affine transformation. The work highlights the sufficiency of co-modular grid cells for path integration.

## 2. Results

### 2.1. An internal representation of space can be constructed from grid cell activity

We present a novel algorithm that reconstructs movement trajectories from grid cell activity (Fig. 1). The method proceeds in two stages. First, using persistent cohomology and toroidal coordinates, we assign toroidal coordinates to each population vector. This constructs a path on the grid cell torus as the subject moves (Fig. 1A-D). Second, we “lift” this path on the torus to the plane, thereby reconstructing the subject’s movement in physical space (Fig. 1E).

**Figure 1.**
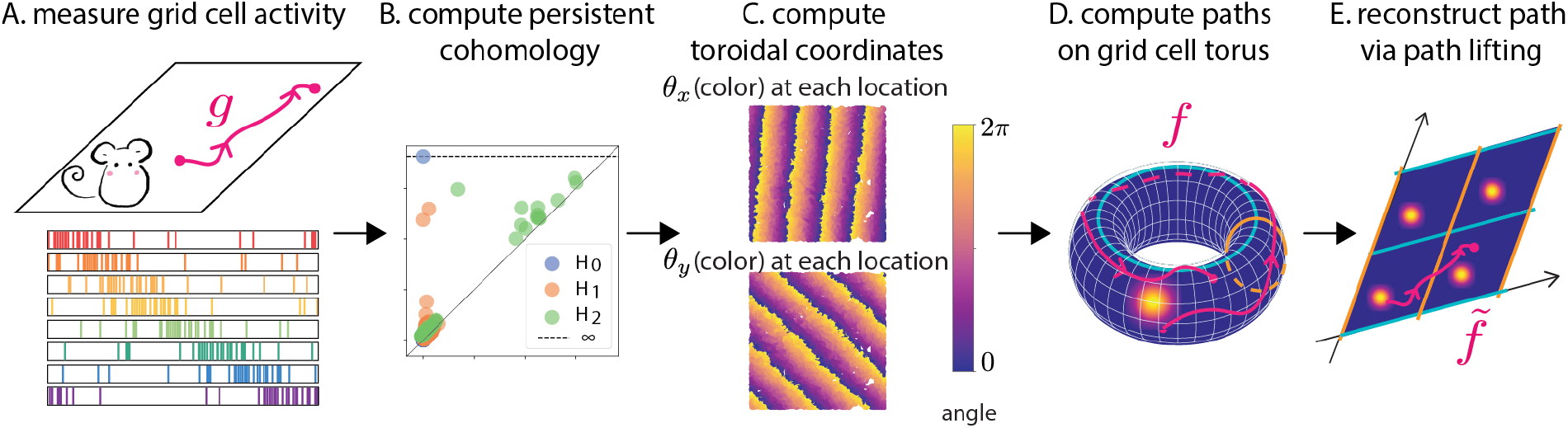
Constructing an internal representation of space from grid cell activity. **A**. The input data is grid cell activity collected while the mouse moves in an environment. Grid cell population activity is represented as a population vector *P*(*t*) evolving over time. **B**. Persistent cohomology indicates that the population vectors are organized on a torus. **C**. Each population vector *P*(*t*) is assigned toroidal coordinates 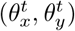. Here, if the mouse is at location (*x, y*) at time *t*, we show the toroidal coordinates 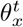 (top) and 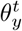 (bottom) by color values over location (*x, y*). **D**. The toroidal coordinates form a path *f* on the grid cell torus. **E**. We finally lift *f* to a path 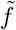 in ℝ^2^ that matches the subject’s movement up to an affine transformation.

#### From grid cell activity to path on a torus

The input is grid cell activity from a subject navigating a spatial environment, represented as a *G* × *T* matrix *A*, where *G* is the number of grid cells and *T* is the number of time bins ^2^. The (*i, j*)^th^ entry represents the activity of neuron *i* at time bin *j*^3^. Each column of *A* corresponds to the population vector *P*(*t*) at time *t*. Although these vectors live in *G*-dimensions, the collection 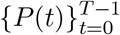 resides on a low-dimensional manifold called a torus [20]. To confirm this, we construct a Vietoris-Rips filtration, which is a nested sequence of simplicial complexes built by connecting population vectors whose dissimilarity falls below an increasing threshold (see *Materials and Methods*). Persistent (co)homology, computed on this filtration, confirms that this low-dimensional manifold has the homology of a torus: one connected component, two 1-dimensional cycles, and one 2-dimensional void (see Fig. 1B). We refer to this manifold as the *grid cell torus*.

Each population vector *P*(*t*) is then parametrized by toroidal coordinates [34, 35] (*Materials and Methods*), reflecting its position on the grid cell torus. Formally, we define the map Θ : *{*0, 1, …, *T* − 1*}* → *S*^1^ *× S*^1^ by

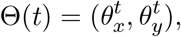

where 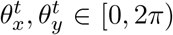. Here, *S*^1^ denotes a circle, and the torus is represented by a product of two circles, *S*^1^ *× S*^1^. Each 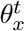 and 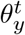 represents angles on each circle. See Figure 1C.

The sequence 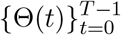 forms a (discrete) path on the grid cell torus (Fig. 1D).

#### From path on grid cell torus to a path in the plane

Once the path on the torus is obtained, we finally recover the movement trajectory in ℝ^2^. Conceptually, we “unwrap” the path on the torus into ℝ^2^, which we accomplish via path liftings to covering spaces [29]. Given a covering map *p* : ℝ^2^ → *S*^1^ *× S*^1^ and a continuous path *f* : [0, *T* − 1] → *S*^1^ *× S*^1^ defined on the interval [0, *T* − 1], one can lift the path *f* to 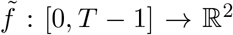 so that 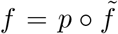, i.e., the following diagram commutes (see SI Section 1.1 for details).

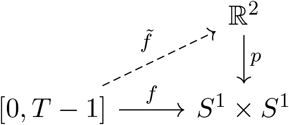

Conceptually, the map *p* folds ℝ^2^ into a square torus by tiling the plane into parallelograms and mapping each parallelogram to one copy of the square torus (see SI Fig. 2). In an idealized setting where grid cell responses at the same physical location are identical, the toroidal coordinates 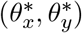 will be identical at every time point *t* at which the subject visits location (*x, y*). In such idealized settings, the map *p* is defined as 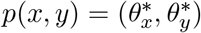.

**Figure 2.**
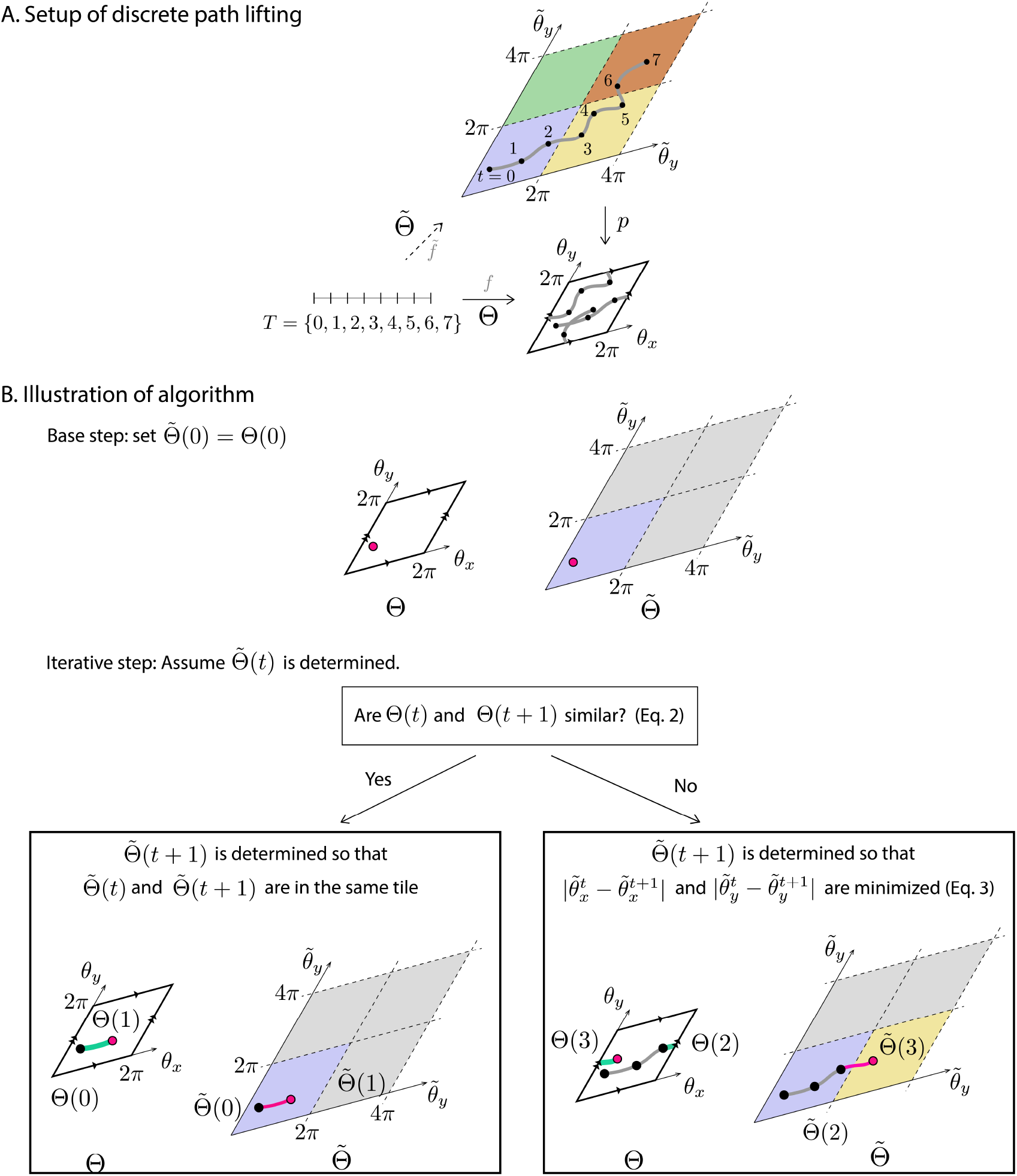
Lifting a discrete path Θ on the torus to a path 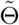 in ℝ^2^. **A**. Setup: given a discrete path Θ : *{*0, 1, …, *T* − 1*}* → *S*^1^ *× S*^1^ on the torus, and the goal is to construct a lifted path 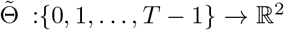 such that 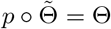, where *p* : ℝ^2^ → *S*^1^ *× S*^1^ is a covering map. **B**. Algorithm. *Base step:* 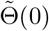 is placed in the tile closest to the origin (blue). *Iterative step:* Given 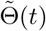, the next lift 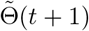 is determined by comparing the consecutive toroidal coordinates Θ(*t*) and Θ(*t* + 1) via Eq. 2. If they are similar (“Yes” branch), the underlying path (green) is assumed to not cross a torus edge and 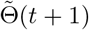 is placed in the same tile as 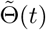. Otherwise (“No” branch), the underlying path (green) is assumed to cross at least one edge and 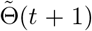 is placed in an adjacent tile, chosen to minimize 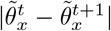 and 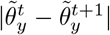(Eq. 3).

Here, *f* denotes a continuous path on the grid cell torus traced by the population vectors throughout the experiment. We treat the sequence 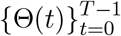 as samples of the path *f*. The goal is to create a discrete path 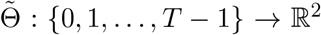 such that the sequence 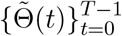 are samples of 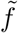. In particular, 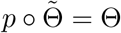. See Fig. 2A for a visualization of this setup.

We define 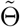 by lifting segments of the discrete path 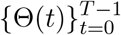 to various parallelograms of ℝ^2^. This will be done by defining 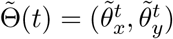 via

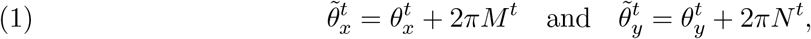

where *M* ^*t*^ and *N* ^*t*^ are integers specifying the tile in which the lifted point inhabits.

The algorithm inductively determines *M* ^*t*^ and *N* ^*t*^. Conceptually, we define 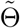 so that if two consecutive time points *t* and *t* + 1 have similar toroidal coordinates Θ(*t*) and Θ(*t* + 1), then their lifts 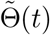 and 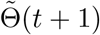 live in the same tile. Otherwise, the lifted points live in different tiles. We say that the toroidal coordinates are similar if

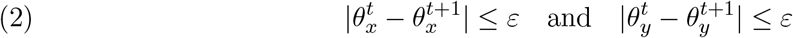

for some proximity threshold *ε*. The proximity threshold *ε* controls which consecutive time points are tested for nontrivial lifts: if the coordinate difference exceeds *ε*, the algorithm evaluates whether the path has crossed a torus edge; otherwise, the two points are lifted to the same tile. Importantly, *ε* only flags candidates – whether a nontrivial lift actually occurs is decided by a distance comparison in Equation 3. We select *ε* from the distribution of consecutive co-ordinate differences, which concentrate near 0 and 2*π*, by choosing a value that lies below the cluster near 2*π*. This ensures that nearly all potential edge crossings are tested. (See *Materials and Methods* and Fig. 9 for details). The path reconstruction is largely insensitive to the precise choice of *ε* (SI Fig. 9).

We now describe the algorithm that defines *M* ^*t*^ and *N* ^*t*^ from Equation 1 inductively. First, we choose the tile closest to the origin (Fig. 2, base step) and define 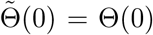 by setting *M* ^0^ = *N* ^0^ = 0.

For each pair of consecutive time points *t* and *t* + 1, we test if the toroidal coordinates Θ(*t*) and Θ(*t* + 1) are similar via Equation 2. If the two coordinates are similar, then Θ(*t* + 1) is lifted to the same tile as 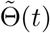 by setting *M* ^*t*+1^ = *M* ^*t*^ and *N* ^*t*+1^ = *N* ^*t*^.

If Θ(*t*) and Θ(*t* + 1) fail to satisfy Equation 2, there are two possibilities for the underlying path *f* |_[*t,t*+1]_. The first possibility is that the path *f* |_[*t,t*+1]_ did not cross any of the edges of the torus (Fig. 2, “Yes” branch, green path), in which case 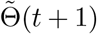 should remain in the same tile as 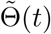. We set *M* ^*t*+1^ = *M* ^*t*^ and *N* ^*t*+1^ = *N* ^*t*^.

The second possibility is that *f* |_[*t,t*+1]_ crossed at least one edge of the square torus (Fig. 2, “No” branch, green path segment), in which case 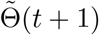 should lie in a tile adjacent to that of 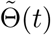. This amounts to setting *M* ^*t*+1^ = *M* ^*t*^ ± 1 and (or) *N* ^*t*+1^ = *N* ^*t*^ ± 1 ^4^.

In practice, the underlying path *f* is unobserved, so we do not know which case applies. We therefore compare both cases and choose the lift that results in a smaller distance to 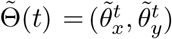 for each coordinate. That is,

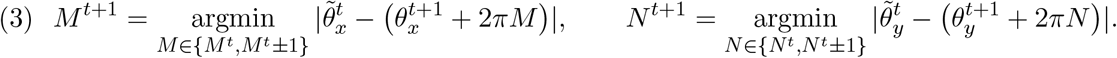

We then use *M* ^*t*+1^ and *N* ^*t*+1^ to define 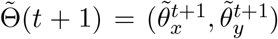 via Equation 1. See Fig. 2 for an illustration of the algorithm. We repeat this process for all time points to obtain the full lifted sequence 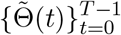. We consider 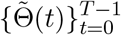 as the reconstructed movement path in ℝ^2^ (Fig. 2A). In the following sections, we show that this reconstructed path closely matches the subject’s true trajectory up to an affine transformation. We emphasize that the subject’s physical location is not used in the path reconstruction process.

#### Measuring the quality of the path reconstruction

We evaluate reconstruction quality at two scales. The global reconstruction error measures the fidelity of the entire reconstructed trajectory, computed as the mean Euclidean distance between the original and reconstructed paths after optimal affine alignment, normalized by the size of the environment (see *Materials and Methods*). Local reconstruction error, on the other hand, captures whether the reconstruction preserves local geometry even when the global shape may be distorted. To compute local reconstruction error, the original movement path and the reconstructed path are first split into shorter path segments. For each segment, we align the corresponding movement segment and reconstructed path segment and compute the normalized, mean Euclidean distance as described in *Materials and Methods*. As we will show, the distinction between these two scales is important: a reconstruction can be locally faithful while the global geometry is distorted (see Fig. 7), possibly because local reconstruction can remain accurate even when small lifting errors accumulate and distort the global path (see SI Section 4.3).

### 2.2. Grid cell activity accurately reflects the geometry of the environment

A fundamental question is whether the proposed method can faithfully reconstruct the true movement trajectory from grid cell activity. To test this, we first consider simulated grid cell activity where the ground-truth trajectory is known.

We simulated mouse trajectories of length 25, 000 in three environments with zero, one, and two holes (Fig. 4A, *Materials and Methods*). Grid cell activity was generated using a continuous attractor network (CAN) model [5, 20], implemented on a 56 × 44 grid cell network with a shared spatial resolution. This produced simulated activity of *G* = 2, 464 grid cells over approximately *T* = 599, 999 time bins (see *Materials and Methods*). We then applied the proposed pipeline to reconstruct the movement paths (Fig. 3). A visualization of the toroidal coordinates over the physical environment reveals that multiple locations elicit similar toroidal coordinates, indicating that path reconstruction will require nontrivial lifts (Fig. 4B).

**Figure 3.**
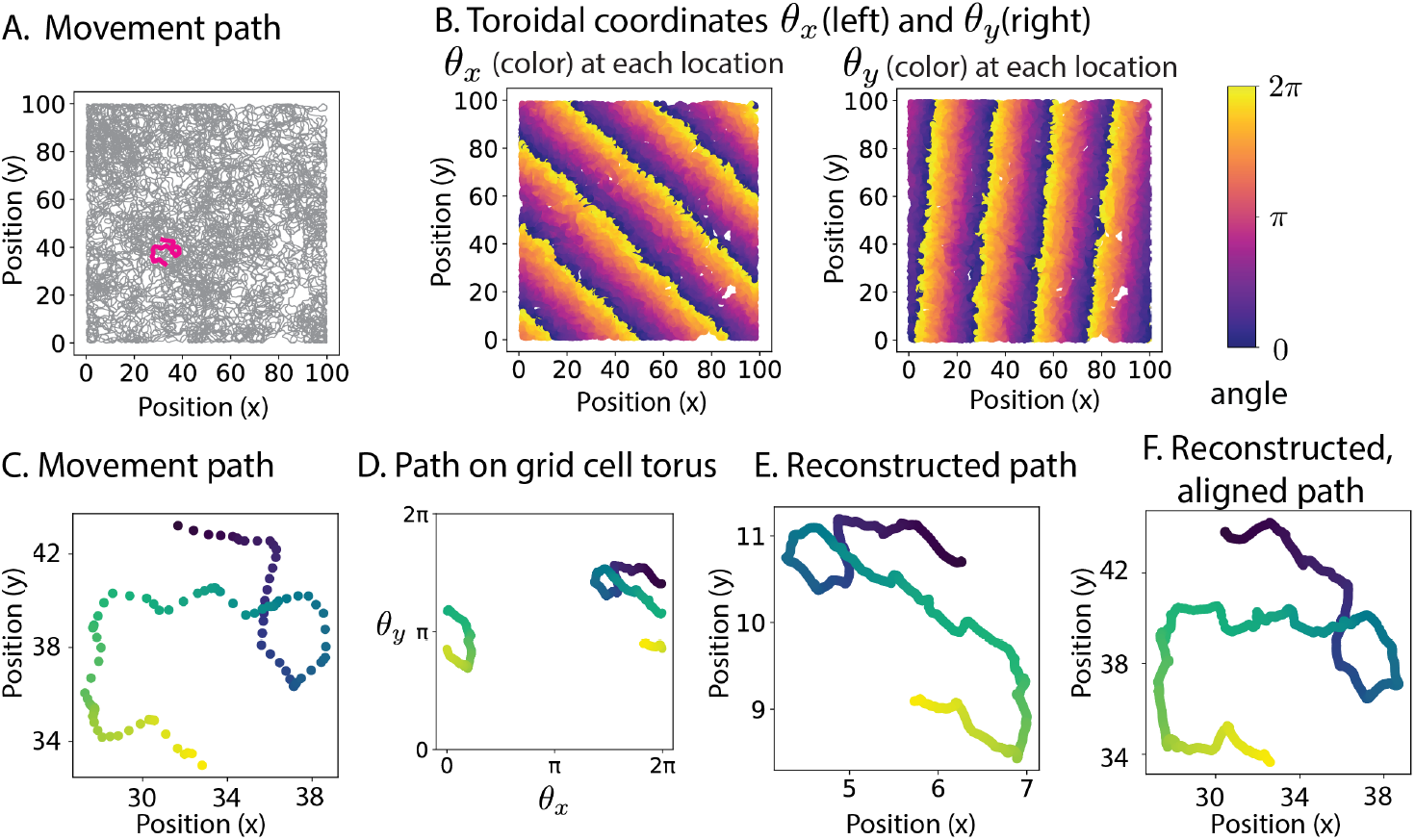
Illustration of path lifting on a simulated CAN data (56 × 44 grid cell network, *T* = 599, 999 time bins.). **A**. A simulated movement path, with a highlighted segment. **B**. Toroidal coordinates for each location on the map. The repeated values indicate that the map is large enough to require nontrivial lifting during path reconstruction. **C**. Enlarged view of the highlighted segment. The color indicates that the simulated mouse moves from dark to light. **D**. The toroidal coordinates corresponding to the path segment in panel C. **E**. The output of the reconstruction algorithm resembles the original path in panel C. **F**. The reconstructed path, post affine transformation, recovers the original movement path in panel C.

**Figure 4.**
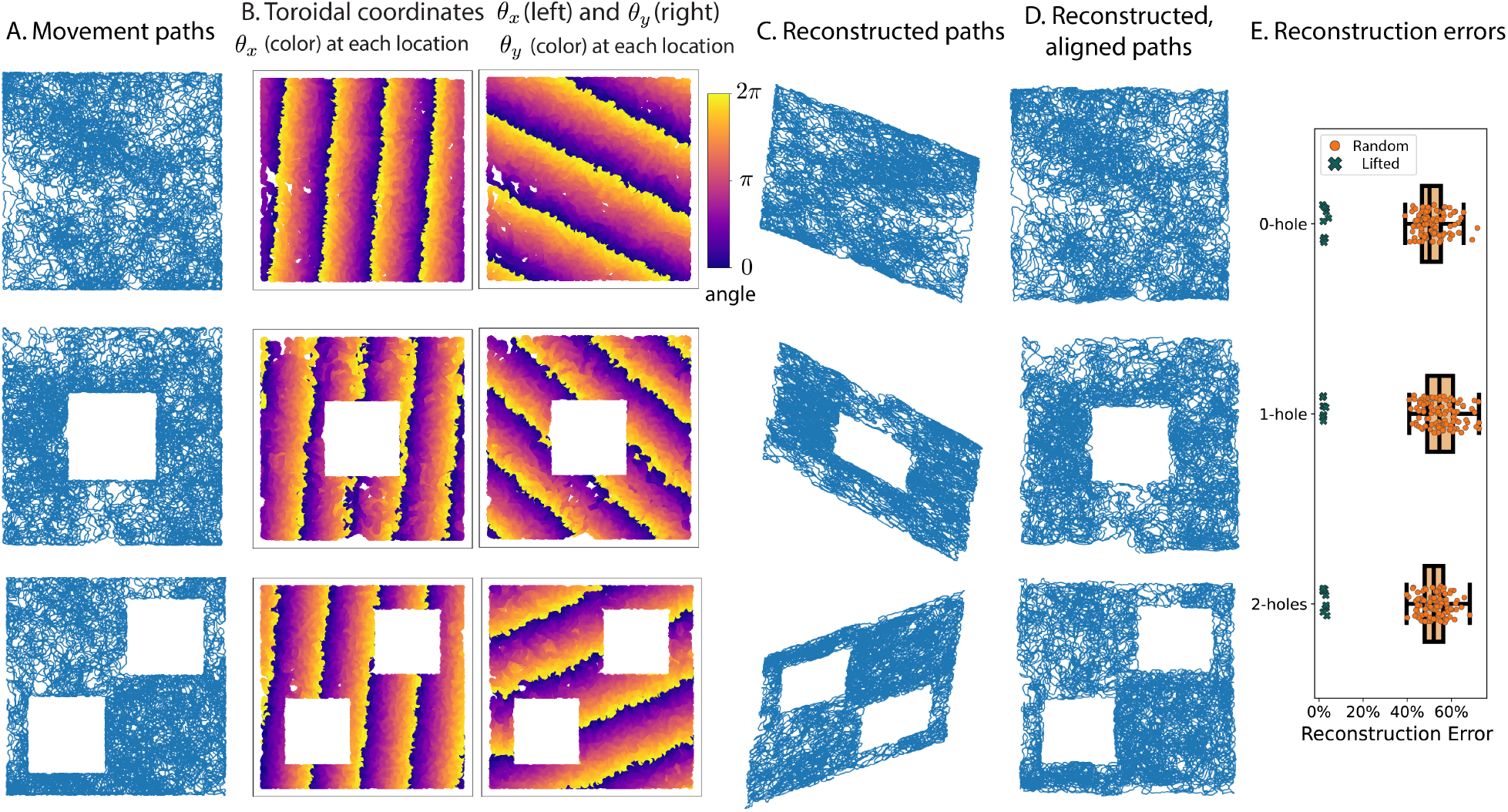
Path lifting on CAN-simulated grid cell activity (*G* = 2, 464, *T* = 599, 999 time bins per simulation.) reconstructs the original movement path. **A**. Simulated movement trajectories in environments with 0, 1, and 2 holes. **B**. Toroidal coordinates for each location on the map. **C**. Reconstructed paths from the simulation of mouse movement on maps with 0, 1, and 2 holes reflect the topology of the maps. **D**. After optimal affine alignment, the reconstructed paths resemble the original movements in panel A. **E**. Reconstruction errors across 10 independent trials. For each environment, the error between simulated movement paths and reconstructed paths (teal) are compared against random baseline (orange), computed as the error between pairs of independently simulated trajectories in the same environment. The reconstruction errors are significantly smaller than the random baselines in all three environments.

The reconstructed trajectories preserved the global topology of the environment. Visualizations of the reconstructed paths captured the presence of holes in the environment (Fig. 4C). Indeed, the persistence diagrams computed from the reconstructed paths revealed the correct number of connected components and one-dimensional holes in the physical map (SI Fig. 20C). These results demonstrate that activity of a single grid cell module provides sufficient information to recover the topological features of the environment.

We then investigated how well the reconstructed path resembles the geometry of the original movement path. We first aligned the reconstructed path to the ground-truth trajectory via an optimal affine transformation (*Materials and Methods*). In each simulation, the aligned reconstructed path closely matched the original, not only preserving the holes but also reflecting spatial patterns such as frequently visited regions (Fig. 4D). Quantitatively, the mean reconstruction error across 10 simulations (Fig. 4E, teal) was significantly lower than the errors from random reconstructions (Fig. 4E, orange) (z-scores: −7.6, −5.6, −8.4), where random reconstruction error refers to the error between two randomly simulated paths in the same environment (see *Materials and Methods*). The local reconstruction errors were computed using paths of length 10,000 time bins, resulting in 59 local paths. The mean and standard deviation of local reconstruction errors for the 1-hole environment were 1.2 ± 0.3%. See SI Fig. 20E for example local paths and their reconstructions.

We next tested robustness to noise in neural activity. To mimic spontaneous firing, we added Gaussian-shaped noise events of peak height *h* and variance *σ*^2^ at random time points to the simulated grid cell activity. The number of noise events added was controlled by a proportion parameter *p* (*Materials and Methods*).

When spontaneous activity is introduced to a relatively small proportion of time points, for example, *p <* 10%, the reconstructions remained highly accurate even under large noise variance *σ*^2^, with errors comparable to those from the original data (see Fig. 5 and Table 1). Even when the resulting activity traces look visibly very different from the original, the reconstruction errors remained small (see Fig. 5 and Table 1). However, at higher *p*, the toroidal structure of the grid cell activity degraded, leading to poor or failed reconstructions (see Table 1 and SI Fig. 4). These results show that the reconstruction method is robust to moderate levels of noise but fails when spurious activity overwhelms the toroidal organization of grid cells. See SI Section 3 for a more thorough analysis of the robustness of the method against various kinds of noise in neural activity, including spontaneous firing, neural activity suppression, and temporal shifts.

**Figure 5.**
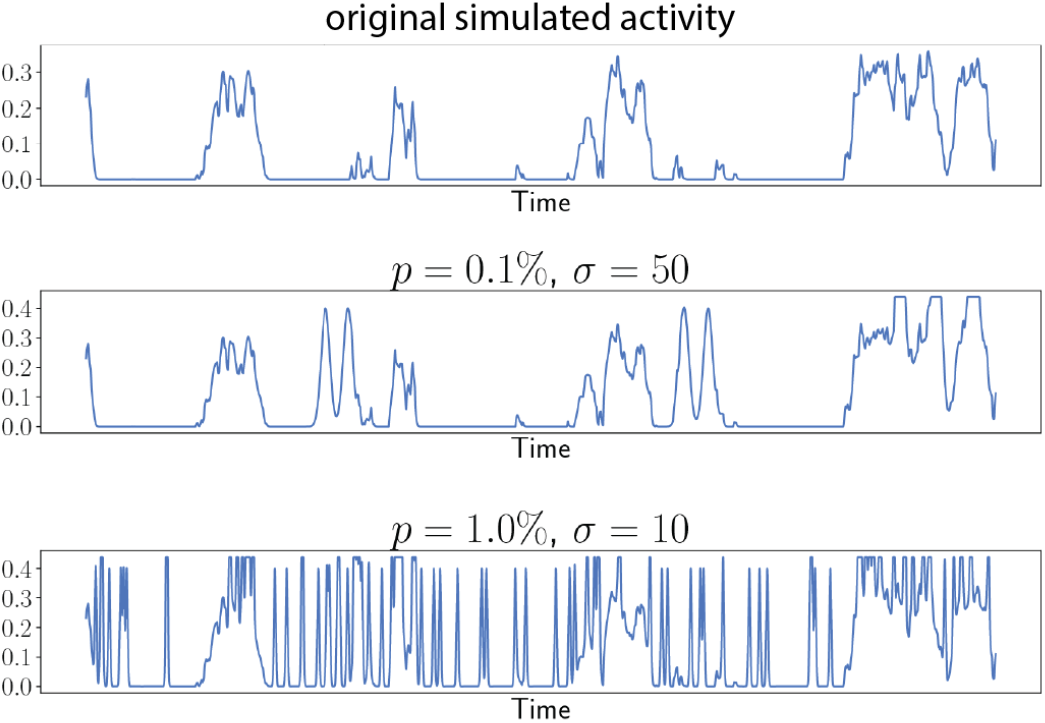
Example simulated grid cell activity with spontaneous firings that lead to low path reconstruction errors. (Top) Example activity trace from a CAN simulation. (Center) Simulated activity with additional spontaneous activity, generated with *h* = 0.4, *p* = 0.1% and *σ* = 50. The mean global reconstruction error for such noisy activity is 2.115% (see Table 1). (Bottom) Activity trace with additional spontaneous activity, generated with *h* = 0.4, *p* = 1% and *σ* = 10. The mean reconstruction error is 4.894% (see Table 1).

**Table 1.** Average reconstruction errors (%) between the original trajectory and the reconstructed paths over 10 trials. Here, the maximum height of the spontaneous firing is fixed at *h* = 0.4. The rows represent the proportion of times during which a grid cell randomly fired, and the columns represent the variance *σ* of the noise added. An entry of N/A indicates that the method failed to compute toroidal coordinates in all 10 trials.

| | | Standard Deviation ($\sigma$) | | | |
| --- | --- | --- | --- | --- | --- |
|  |  | 1 | 10 | 50 | 100 |
| Proportion ($p$) | 0.1 % | $1.647 \pm 0.0295$ | $1.723 \pm 0.0413$ | $2.115 \pm 0.1768$ | $49.480 \pm 32.2798$ |
| | 0.5 % | $1.683 \pm 0.0327$ | $1.903 \pm 0.0571$ | $42.327 \pm 2.2691$ | $44.736 \pm 3.7464$ |
| | 1 % | $1.695 \pm 0.0492$ | $4.894 \pm 7.3985$ | $41.785 \pm 2.0993$ | $45.528 \pm 5.9802$ |
| | 5 % | $1.819 \pm 0.0922$ | $44.294 \pm 4.6828$ | N/A | N/A |
| | 10 % | $14.744 \pm 15.2993$ | $44.176 \pm 5.4732$ | N/A | N/A |

### 2.3. Path reconstructions recovers one-dimensional environment from grid cell activity

We applied the method to experimental data, asking whether one-dimensional trajectories can be reconstructed from real grid cell recordings. We analyzed a publicly available dataset of grid cell activity in mice navigating a 320 cm virtual build-up track [44]. Here, whenever the mouse reached the end of a track, the mouse was teleported to the start without visual discontinuity (Fig. 6A). We analyzed 441 such runs on the linear track. The co-modular grid cells were identified via clustering on spectrograms (see [44] for details). The dataset provides firing rates that were preprocessed by the original authors as follows: spikes were binned into 2-cm spatial bins along the track, divided by the time occupancy per bin, and smoothed with a 2-bin Gaussian kernel. Each continuous run on the 320 cm track thus produced grid cell firing rates over 160 spatial bins.

**Figure 6.**
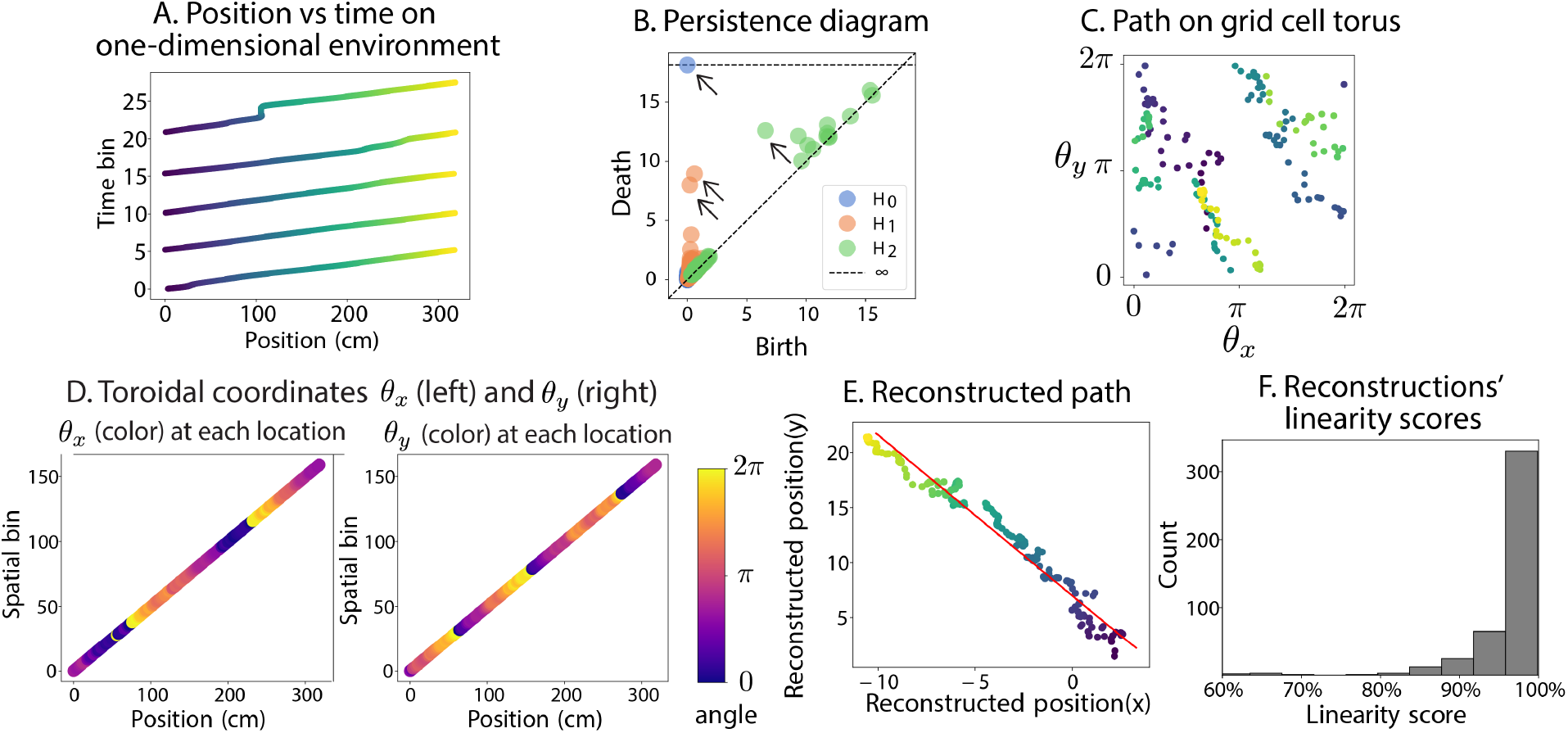
Path reconstruction recovers one-dimensional environment from grid cell activity. Data are from mouse *N* 2 (dataset “N2 200203 buildup track”; [44]) navigating a 320 cm virtual build-up track. 44 co-modular grid cells were identified. Firing rates were provided in 2*cm* spatial bins (160 bins per run, 441 total runs). **A**. Mouse position over time across 5 runs; each rising segment corresponds to one traversal of the track, after which the mouse is teleported to the start. A single run is highlighted in pink. **B**. The persistence diagram confirms that grid cells are organized on a torus: one connected component (*H*_0_), two one-dimensional cycles (*H*_1_), and one two-dimensional void (*H*_2_). **C**. Example path on the grid cell torus corresponding to a single run. For each time point *t*, the corresponding toroidal coordinates *θ*_*x*_ and *θ*_*y*_ are plotted. **D**. Toroidal coordinates from panel C visualized over position. Because the firing rate data is provided in 2*cm* spatial bins, the toroidal coordinates are also computed for each spatial bin. Each point on the plot corresponds to one spatial bin in a fixed run, plotted at its track position (*x*-axis) and spatial bin index (*y*-axis). Color encodes the toroidal coordinates *θ*_*x*_ (left) and *θ*_*y*_(right). **E**. The reconstructed path lies close to a one-dimensional line. The red line indicates the line spanned by the first principal component (PC1) of PCA. **F**. Distribution of linearity scores (variance explained by PC1) across 441 runs; median = 98.8%.

From 44 identified grid cells, the persistence diagrams confirmed a toroidal organization of population activity (Fig. 6B). While toroidal coordinates were computed from the 44 grid cells over the full duration of the experiment, path reconstructions were performed separately on each of the 441 continuous runs of the track. For each run, the toroidal coordinates (*θ*_*x*_, *θ*_*y*_) cross the edges of the grid cell torus (Fig. 6C,D), indicating that the path reconstruction will involve nontrivial liftings. The reconstructed paths recovered the one-dimensional structure of the environment (see Fig. 6E). The one-dimensional nature of a reconstructed path was quantified by the variance explained by the first principal component, which we call “linearity score.” Across all 441 runs, the median linearity score was 98.8% (Fig. 6F). For comparison, the median linearity score of 59 local paths from a simulated trajectory in the 0-hole world (see SI Fig. 20E for examples) was 79.2%. This analysis demonstrates that one-dimensional spatial structure can be reliably decoded from experimental grid cell activity.

### 2.4. Local paths can be reconstructed from grid cell activity in a two-dimensional environment

Finally, we tested the method on two-dimensional experimental data that is publicly shared in [20]. Extracellular spikes from grid cells in layers II and III of MEC-parasubiculum were recorded while rats explored a square open-field arena of size 1.5*m* × 1.5*m* (Fig. 7A). Six grid modules were identified by clustering. Here, we report the analysis on one of the datasets (rat *R*, module 1, day 2, OF), which consisted of 111 co-modular grid cells recorded over 21.1 minutes (*T* = 126, 728 time bins, 1 time bin = 10ms) of open-field foraging.

**Figure 7.**
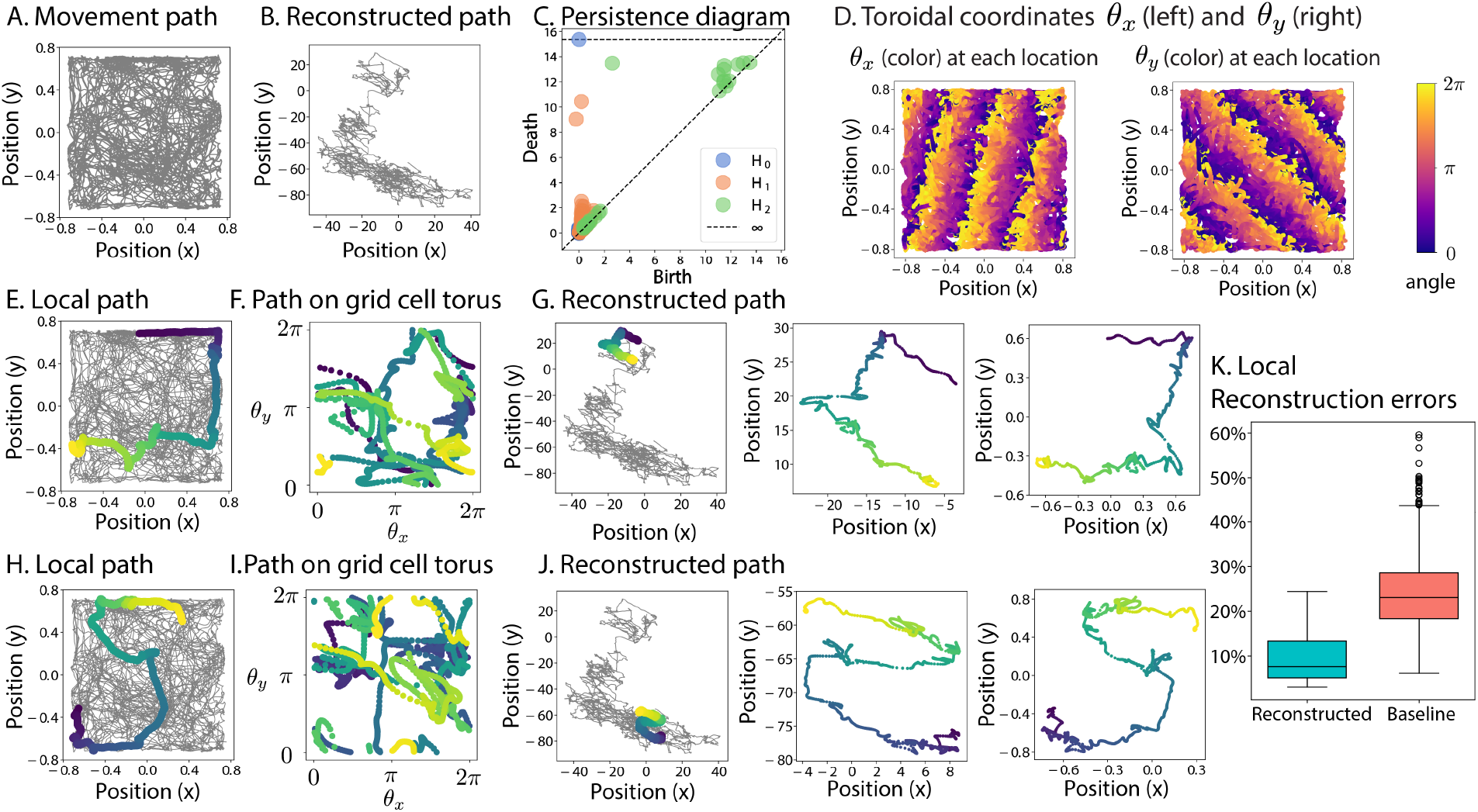
Reconstruction of local paths from two-dimensional experimental data [20] (rat *R*, module 1, day 2, open-field session; 111 co-modular grid cells). **A**. The original trajectory of a rat exploring a 1.5*m* × 1.5*m* open-field arena. **B**. The reconstructed global trajectory, which differs in overall shape from the original path. **C**. The persistence diagram indicates that the grid cells are organized on a torus. **D**. A visualization of the toroidal coordinates for each location. **E**. An example local path. **F**. The toroidal coordinates corresponding to panel E involve non-trivial liftings. **G**. A highlight of the reconstructed segment in panel B (left), the reconstructed path, before affine transformation (center), and after affine transformation (right). **H - J**. Another example local path and its reconstruction. **K**. Distribution of local reconstruction errors: pairs of original local paths and reconstructed paths (left) show significantly smaller errors than baseline consisting of mismatched local paths (right) (*t*(2014) = −14.6, *p <* 0.0001).

Globally, the reconstructed path differed in overall shape from the true trajectory (Fig. 7B). However, when the analysis was restricted to shorter local paths corresponding to 20-second intervals, the reconstructed paths were highly consistent with the original movement paths (Fig. 7E-J). For example, the reconstructed paths in Fig. 7G and J resemble the geometry of the original local paths in Fig. 7E and H, despite requiring many non-trivial lifts across torus edges (Fig. 7F, I).

To quantify the quality of local path reconstructions, we compared the local reconstruction errors between the original and lifted path segments against a baseline distribution of mismatched segment pairs. Local reconstructions had significantly lower errors (mean 9.5%) than the null baseline (mean 23.7%, s.d. 7.6%, Fig. 7K), confirmed by an independent t-test (*t*(2014) = − 14.6, *p <* 0.0001). These results establish that while global reconstructions may deviate from the true path, local trajectories can be faithfully recovered.

Several factors can contribute to the discrepancy between local and global path reconstructions. First, noise in the toroidal coordinates can distort the lifted path (compare Fig. 7D to Fig. 3B). Second, when the sampled time points are too sparse, two types of lifting errors can occur: toroidal coordinates Θ(*t*) and Θ(*t* + 1) that should be lifted to the same tile may be lifted to different tiles (see Fig. 8A), or coordinates Θ(*t*) and Θ(*t* + 1) that should be lifted to distinct tiles may end up lifting to the same tile (see Fig. 8B). Accumulated errors of this type can cause large-scale distortions in the global path reconstruction (see SI Section 4.3).

**Figure 8.**
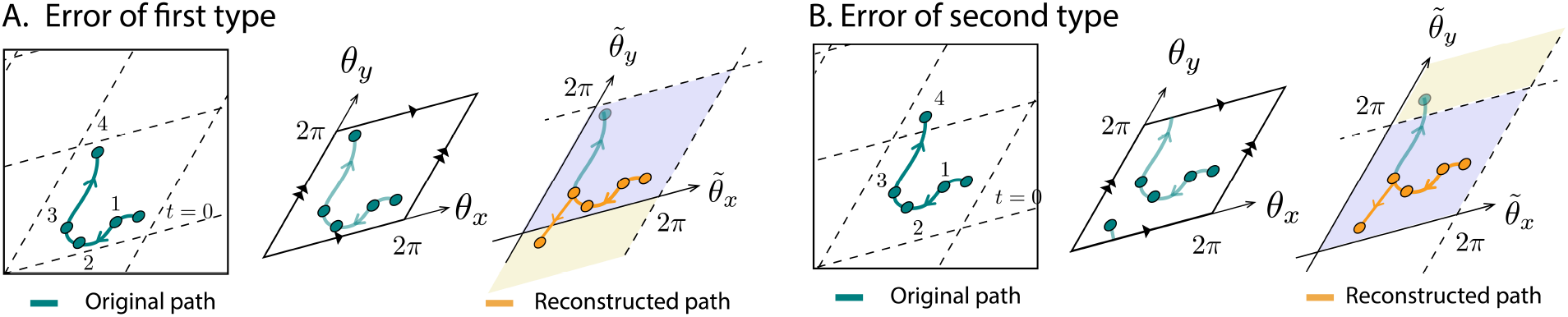
Two possible errors in path reconstruction arising from sparsity of time points. **A**. The first type of error occurs when two consecutive toroidal coordinates are lifted to two distinct tiles when they should be lifted to a single tile. (Left) Original movement path. Circles indicate the location at select time points. (Center) The corresponding toroidal coordinates. (Right) The algorithm lifts the toroidal coordinates Θ(0), …, Θ(3) to the blue tile. Because 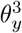 and 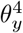 are dissimilar, 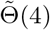 is in a different tile, shown in yellow. The resulting reconstructed path (orange) deviates from the original path (green). **B**. The second type of error occurs when two consecutive toroidal coordinates are lifted to the same tile when they should be lifted to different tiles. Here, the toroidal coordinates 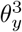 and 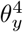 have a small enough difference so the algorithm lifts Θ(3) and Θ(4) to the same tile. Again, the reconstructed path (orange) deviates from the original (green).

## 3. Discussion

In this study, we introduced and validated a topological framework for decoding spatial trajectories from grid cell population activity. By identifying toroidal coordinates through persistent cohomology and lifting paths from the torus to the plane, we effectively reconstruct movement trajectories up to an affine transformation. This approach decodes spatial trajectories without access to external positional cues or knowledge of grid phases, offering a new computational perspective on spatial representation of the grid cell system. Our results show that the toroidal organization of grid cell activity is not merely a descriptive feature but can be functionally leveraged to recover spatial information.

The proposed method builds on the work of Gardner et al [20], who used persistent cohomology to reveal the toroidal structure of grid cell activity and assigned toroidal coordinates to population vectors using the circular coordinates construction of de Silva et al [35]. While prior work characterizes the structure of the neural manifold, present work shows that these toroidal coordinates can be used to decode spatial trajectories from it. The key methodological contribution is path lifting – unwrapping paths on the grid cell torus into paths in the plane via covering maps.

The proposed method complements previous decoding approaches in that it only requires the activity of a single grid module (as opposed to multiple grid modules) [26, 38] and that it doesn’t involve any training process. Furthermore, the method doesn’t require any phase information of the grid modules, enhancing the applicability of the method.

Recent work has shown that individual grid cell spike trains are often too irregular to convey the lattice structure of the firing fields, and that extracting lattice structure from spike trains requires parameter regimes in which the animal’s trajectory passes through neighboring grid fields in sequence without omissions [11]. In contrast, our method operates on population vectors – the simultaneous activity of co-modular grid cells at each time step. The collection of these population vectors forms a torus, which is then used for path reconstruction. Because we analyze the population-level activity, we do not require any individual cell to exhibit regular firing activity.

In order for the method to reconstruct paths accurately, several assumptions must be met. The presented method requires that the grid cells are recorded for a sufficiently long time so that the toroidal structure of the population vectors becomes clear. Furthermore, accurate path reconstruction depends on sufficiently dense temporal sampling. As discussed in Fig. 8, having insufficient time points can lead to poor lifting. On the other hand, including too many time points can lead to slow computation, especially when computing the toroidal coordinates. In particular, the analysis of two-dimensional experimental data involves a preprocessing that selects the relevant time points. Such preprocessing must be done in a manner that preserves the toroidal structure of grid cells.

The affine transformation used to align the reconstructed path to the ground-truth trajectory serves as an evaluation tool to quantify reconstruction accuracy; it is not part of the decoding algorithm itself. It is unclear whether the brain needs to resolve this affine ambiguity to perform path integration. The path reconstruction in this work can be interpreted as performing path integration over basis vectors of ℝ^2^ that are not necessarily orthogonal. If the brain’s internal coordinate system similarly operates in a non-orthogonal basis, then no affine correction would be required to track relative position.

If, however, alignment with the geometry of the physical environment is needed, the affine ambiguity could in principle be resolved in at least two ways. First, combining information from multiple grid modules with different spacings and orientations is sufficient to recover absolute position, because the ambiguity inherent in any single module’s periodic code is eliminated when multiple modules are combined [38]. Second, other spatially tuned cell populations, including place cells and boundary cells, as well as visual landmarks, could provide an external anchor for mapping from the brain’s toroidal coordinate system to the physical environment. These possibilities are consistent with the broader view that spatial navigation relies on the coordinated activity of multiple cell types.

There are several directions for future research. Incorporating interpolation and probabilistic inference during path lifting could improve robustness to sparse or noisy data. Combining grid modules of different phases may provide a more comprehensive and precise decoding framework. Future research should also examine how the toroidal organization of grid cells integrates with other spatially tuned cell populations such as place cells to support path integration and spatial navigation. Because place cells provide a non-periodic spatial code, they could help disambiguate the choice of lift during path reconstruction from grid cell activity. For example, a place cell that fires at two similar time points would indicate that the lifted coordinates at the corresponding times should occupy a similar region in ℝ^2^, providing a constraint that guides the path reconstruction algorithm. Topological methods have been used to reconstruct stimulus spaces from place cell activity[10], and integrating such information with the present framework is a natural direction for future work.

Beyond neuroscience, the method offers new perspectives on artificial navigation systems. Recent work has shown that grid-like representations emerge in deep networks trained to perform path integration, enabling vector-based navigation in artificial agents [1]. Grid cell-inspired path integration and state estimation have also been explored for mobile robots [36, 47]. Our framework suggests a complementary approach: if an artificial agent’s internal representation exhibits toroidal structure, whether learned or engineered, path lifting could recover spatial trajectories and aid in position estimation without external cues.

## 4. Materials and methods

### 4.1. Simulation of grid cells

#### 4.1.1. Simulated Mice Trajectory

A two-dimensional random walk simulation was developed to model the exploratory behavior of mice within a bounded environment (100 × 100). The environment was defined by spatial boundaries and obstacle parameters, including the sizes and positions of holes. A mouse agent was initially placed at a randomly selected valid location within these boundaries. At each time step, candidate positions were computed within an angular window of ± 75° relative to the current heading. A step size is randomly drawn up to a predefined maximum, and candidate positions are generated by adding a scaled directional vector to the current position. Only positions that remained within the environment and avoided designated holes were accepted. If no valid candidate position was available, the agent remained stationary for that time step and subsequently adopted a new random heading from the full 360° range. Repeated iterations of this process produced a random trajectory that models the exploratory behavior of mice. All simulated trajectories had length 25, 000.

#### 4.1.2. Grid cell simulation with CAN model

The simulated mice trajectories were then used to simulate grid cell activity using a noiseless CAN model with purely inhibitory recurrent connectivity, following [5, 9]. We use the same model and parameters that were used in [20]. There, the CAN model was used to simulate grid cell activity in response to position data. Here, we use it to simulate grid cell activity in response to simulated mice trajectory.

The network consists of a 56 × 44 neuronal sheet with periodic (toroidal) boundary conditions. The simulated random walk trajectory was then provided as input to the model, which computed the speed *v*(*t*) and head direction *θ*(*t*) at each time step. The activity of neuron *i* at time *t* is determined by

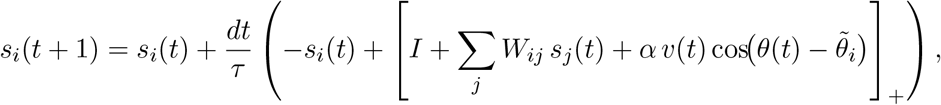

where [*x*]_+_ = max(*x*, 0) is the threshold-linear function, and 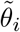 is the preferred direction of neuron *i*. The parameters followed the implementation in [20]: *I* = 1 (constant external input), *α* = 0.15 (velocity modulation), *dt* = 1 (integration time step), and *τ* = 10 (neuronal time constant). *W*_*ij*_ denotes the strength of connection from neuron *j* to neuron *i*, and it was computed as described in [9] with *W*_0_ = −0.02 and connectivity radius *R* = 15.

The activity patterns were initialized by setting 90% of neurons to *s*_*i*_ = 1 (the rest were set to 0) and performing 2, 000 updates to allow the hexagonal bump pattern to stabilize. For computational efficiency, the activity was set to 0 whenever *s*_*i*_ *<* 0.0001.

For each simulation, the result was simulated activity of *G* = 56 × 44 = 2, 464 grid cells over *T* = 599, 999 time bins. The activity values ranged from 0 to approximately 0.45 per time bin, with a mean peak activity of approximately 0.44 per time bin. These amplitudes and time bins are inherent to the model dynamics and should not be interpreted as firing rates in physiological units such as Hz. The grid fields had an average diameter of 14.8 units in an environment of size 100 by 100. See SI Fig. 20D for example firing fields.

We refer the reader to [9] for the original model and [20] for details of the implementation we adapted.

#### 4.1.3. Simulation of grid cells with additional spontaneous activity

To emulate the random firing of grid cells, we incorporated one-dimensional Gaussian noise into the simulated grid cell activity. Given a simulated activity *r*(*t*) with *t* ∈ [0, *T*], we modify *r*(*t*) by adding one-dimensional Gaussian functions *g*_*h,σ*_(*t*) centered at some random value in [0, *T*], with peak height *h* and variance *σ*^2^. Letting *r*_max_ = max_*t*_ *r*(*t*), the maximum value in the simulated data, we construct a noisy activity trace by

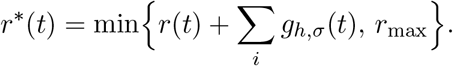

We clip the result so that *r*^∗^(*t*) doesn’t exceed the maximum value *r*_max_ from the original simulation. Here, the number of Gaussian functions added can vary, and the number is determined as some proportion *p* of *T*.

For computational efficiency, a fast approximation routine precomputes a truncated Gaussian curve by identifying the index at which the noise amplitude falls below a specified threshold of 1e-4, thereby limiting the range over which noise is applied.

### 4.2. Topological features of grid cell activity

#### 4.2.1. Persistent cohomology

Persistent (co)homology is a tool in topological data analysis that can be used to identify structural features in neural manifolds. Here, it is used firstly to verify that the grid cells’ population activity is organized in a torus and secondly to compute the toroidal coordinates of each population vector.

Given an activity matrix *A* with *G* grid cells and *T* time bins as input to persistent cohomology computation, the general persistent cohomology computation on the population activity would first create a symmetric *T* × *T* pairwise dissimilarity matrix. However, due to the large size of *T*, persistent cohomology is usually computed on a smaller number of time points *T* ^∗^ (*T* ^∗^ = 250 in simulated data; *T* ^∗^ = 1, 200 in both experimental data, where the selection of *T* ^∗^ time points is described in the following section). Let 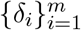 be a collection of nonnegative real numbers satisfying 0 ≤ *δ*_0_ *< δ*_1_ *<* · · · *< δ*_*m*_. Given a parameter *δ*_*i*_, we construct a Vietoris-Rips complex 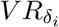 that consists of *T* ^∗^ vertices and has an *n*-simplex [*v*_0_, …, *v*_*n*_] precisely when all pairwise dissimilarity among the listed elements is at most *δ*_*i*_. We then obtain a filtration of Vietoris-Rips complexes

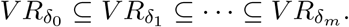

Computing (co)homology in dimensions 0, 1, and 2 with a field coefficient, we obtain a sequence of vector spaces summarizing the connected components, circular features, and voids in each Vietoris-Rips complex. The birth and death of these topological features are summarized in a persistence diagram. A topological feature born at parameter *b* that dies at parameter *d* is represented by a point in the plane with coordinates (*b, d*) (see Fig. 1B). Ripser [3, 42] was used for persistent cohomology computations. For a general introduction to persistent cohomology, we refer the reader to SI Section 1.2 and [7, 13, 14, 21].

For the simulated data, the dissimilarity between two population vectors were computed using Euclidean distance. Persistent cohomology was computed using DREiMac [32], which selects *T* ^∗^ = 250 landmarks for its computation. For the experimental data, where more noise is present, we compute persistent cohomology on *T* ^∗^ = 1, 200 time bins with cosine dissimilarity. The selection of *T* ^∗^ time bins is described below.

#### 4.2.2. Preprocessing and persistent cohomology computation of experimental grid cell activity

When analyzing the one-dimensional and two-dimensional experimental data, we followed the preprocessing pipeline and publicly available code of [20] to construct the grid activity matrix, as described below.

The first step is to obtain a firing rate estimate for each grid cell. For the one-dimensional experimental data [44], the dataset provides firing rates that were spatially binned (2-cm bins) and smoothed by the original authors (see *Data Availability*). For the two-dimensional experimental data, the spike trains were converted to firing rate estimates by convolving with a Gaussian kernel and then sampling, following the preprocessing steps described in [20].

Given *G* co-modular grid cells observed over *T* bins (spatial bins for the one-dimensional experimental data; time bins for the two-dimensional experimental data), let *A* be a *G* × *T* activity matrix, where row *i* of *A* is the activity of neuron *i*. Computing persistent cohomology on large point clouds is computationally expensive and sensitive to outliers, so the population vectors were downsampled and dimension-reduced prior to the computation, following the steps described in [20]. First, 15,000 population vectors with the highest total activity were selected. These were z-scored and projected onto their first six principal components via PCA. The resulting six-dimensional point cloud was further subsampled to *T* ^∗^ = 1, 200 points using a neighborhood-based selection procedure adapted from UMAP, which iteratively selects points with the strongest local neighborhoods. A 1, 200 × 1, 200 symmetric dissimilarity matrix was then computed from these points in the fuzzy topological representation described in [20]. The dissimilarity was computed using cosine dissimilarity. The performance of the path lifting algorithm was largely insensitive to the choice of metric at this stage (see SI Section 4.5). This matrix served as input to the persistent cohomology computation. We refer the reader to the *Methods* section and *Supplementary Methods* of [20] for details.

#### 4.2.3. Circular and toroidal coordinates

Given a point cloud *P* that is arranged in a circular fashion, *circular coordinates* parametrize *P* using a circle-valued map *f* : *P* → *S*^1^, where *S*^1^ denotes a circle. In practice, the range of the circular coordinates will be the angles [0, 2*π*), considered as a circle after identifying 0 and 2*π*. Originally presented in [35], the construction is motivated by the fact that there is a bijection between the equivalence classes of continuous maps from a CW complex *K* to the circle *S*^1^ and the cohomology of *K* with integer coefficients

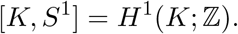

Given a point cloud *P*, persistent cohomology is used to fix a Vietoris-Rips complex *V R*_*δ*_ such that there is a nontrivial circular structure. Then a generator [*η*] ∈ *H*^1^(*V R*_*δ*_; ℤ) is used to assign circular coordinates to each point of *P*. See [35] for details.

Since the torus *S*^1^ *× S*^1^ is a product of two circles, any point on the torus can be parametrized using two circular coordinates as (*θ*_*x*_, *θ*_*y*_), where *θ*_*x*_, *θ*_*y*_ ∈ [0, 2*π*).

On the simulated data, we compute the toroidal coordinates using DREiMac [32], which implements the toroidal coordinates algorithm of [34]. This algorithm extends the circular coordinates construction by decorrelating the two circle-valued maps, ensuring that the resulting toroidal coordinates are geometrically independent.

On the experimental data, we assign toroidal coordinates to each population vector using the cohomological decoding framework described in [20], which adapts the original circular coordinates algorithm [35]. Persistent cohomology and toroidal coordinates were computed for the *T* ^∗^ = 1, 200 selected time points and interpolated to the full set of population vectors. See [20] for details.

We use different methods for computing toroidal coordinates on the simulated and experimental data for the following reason. DREiMac requires fewer preprocessing of neural activity matrix than the decoding framework of [20], making it well-suited for simulated data, where the data involved has less noise. Moreover, it allows us to isolate the effect of individual factors — such as neuron count, temporal resolution, or noise level — on reconstruction quality, without confounding these factors with choices made during the preprocessing of experimental data (see SI Section 4). For the experimental data, however, applying DREiMac directly to the raw population vectors often failed to produce toroidal coordinates (resulting in N/A), due to the higher noise levels and variability inherent in the recordings. We therefore adopted the decoding framework of [20] for the experimental analysis. We note that the two methods yield comparable reconstruction errors when applied to simulated data (SI Fig. 17).

### 4.3. Path lifting

#### 4.3.1. Aligning paths via affine transformation

An affine transformation is a linear mapping that preserves points, straight lines, and planes under rotation, translation, and scaling. Affine transformations of 2-dimensional vectors is defined by a 2 *×* 2 matrix *B* and a vector 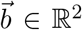. The image of a vector 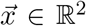 under this transformation is 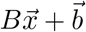.

Given two ordered collections 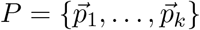 and 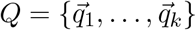 of 2-dimensional vectors, the goal of optimal affine transformation is to find the matrix *B* and 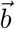 that minimizes the total distance between the transformed source points and their target counterparts. In this work, *P* refers to the (discrete) reconstructed path and *Q* refers to the original movement path. We computed the optimal affine transformation using the `estimateAffine2D` function from the OpenCV package [4], which uses RANSAC (Random Sample Consensus)[17] to iteratively fit candidate transformations to random subsets of three point correspondences and select the transformation that best agrees with the majority of the data. The final transformation is refined using the Levenberg-Marquardt optimization algorithm. We refer to the result of applying the optimal affine transformation to the reconstructed path as the “reconstructed, aligned path.”

#### 4.3.2. Reconstruction error

Let Ψ : {0, 1, … *T* – 1} → ℝ^2^ be the discrete movement path in physical space, where *T* is the number of bins of the grid cell activity (time bins for the simulated and two-dimensional experimental data; spatial bins for the one-dimensional experimental data). Given a reconstructed path 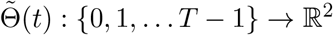, we quantify the dissimilarity between Ψ and 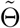 as follows. We first compute the optimal affine transformation aligning 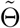 to Ψ as described above, and let 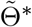 denote the aligned path. The reconstruction error is defined as the mean Euclidean distance between points of Ψ and 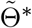, normalized by the environment size:

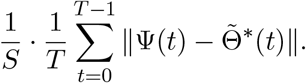

Here, *S* is a normalization factor that allows the error to be interpreted as a fraction of the environment size. When computing the global reconstruction error, we use *S* = 100 (the length and width of the map) for the simulation study and *S* = 1.5 (the length and width of the physical environment in m) for the two-dimensional experimental study. When computing local reconstruction errors of shorter path segments, we use *S* to be the larger of the length and width of the path segment. A lower reconstruction error indicates closer agreement between the original and the reconstructed path.

Throughout this paper, we compare the global and local reconstruction errors against baseline errors computed between pairs of independent movement trajectories in the same environment (Fig. 4E, Fig. 7K). Given two discrete paths Ψ : 0, *{*1, … *T* – 1} → ℝ^2^ and Ω : {0, 1, … *T* − 1}→ ℝ^2^ of equal length, the error between two paths is computed analogously:

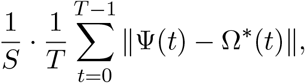

where Ω^∗^ is the transformed path of Ω after aligning it to Ψ. When computing the baseline global reconstruction errors from independent simulations (Fig. 4E), we use *S* = 100. When computing the baseline local errors between two random local paths (Fig. 7K), we set *S* to be the mean size of two paths, where the size of a path is the maximum of the length and width.

#### 4.3.3. Proximity parameter selection

The parameter *ε* determines which consecutive toroidal coordinates are tested for nontrivial lifts. Specifically, if two toroidal coordinates Θ(*t*) and Θ(*t* + 1) satisfy Equation 2, then, they are lifted to the same tile; otherwise, the algorithm evaluates whether the toroidal coordinates should be lifted to adjacent tiles using Equation 3.

A large *ε* causes more consecutive coordinates to be classified as similar, reducing the number of time points tested for nontrivial lifts. A small *ε* increases the number of time points tested. However, because the parameter only identifies candidates for nontrivial lifts – whether a lift actually occurs depends on the distance comparison in Equation 3 – a smaller *ε* generally improves reconstruction accuracy without significant drawbacks. An analysis of the impact of *ε* on the reconstruction error shows that the performance of the path reconstruction is stable across a wide range of *ε* parameters (SI Fig. 9).

To select *ε*, we examine the distribution of maximal coordinate differences 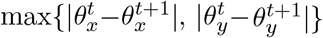 across all consecutive time points. In both simulated (Fig. 9A) and experimental data (SI Fig. 9B, left), these differences concentrate near 0 (reflecting small local movements) and 2*π* (reflecting crossing a torus edge), with the vast majority near 0. Differences near 2*π* correspond to time points at which the path crosses a torus edge and requires a nontrivial lift. We therefore choose *ε* to be smaller than the cluster of values near 2*π*.

**Figure 9.**
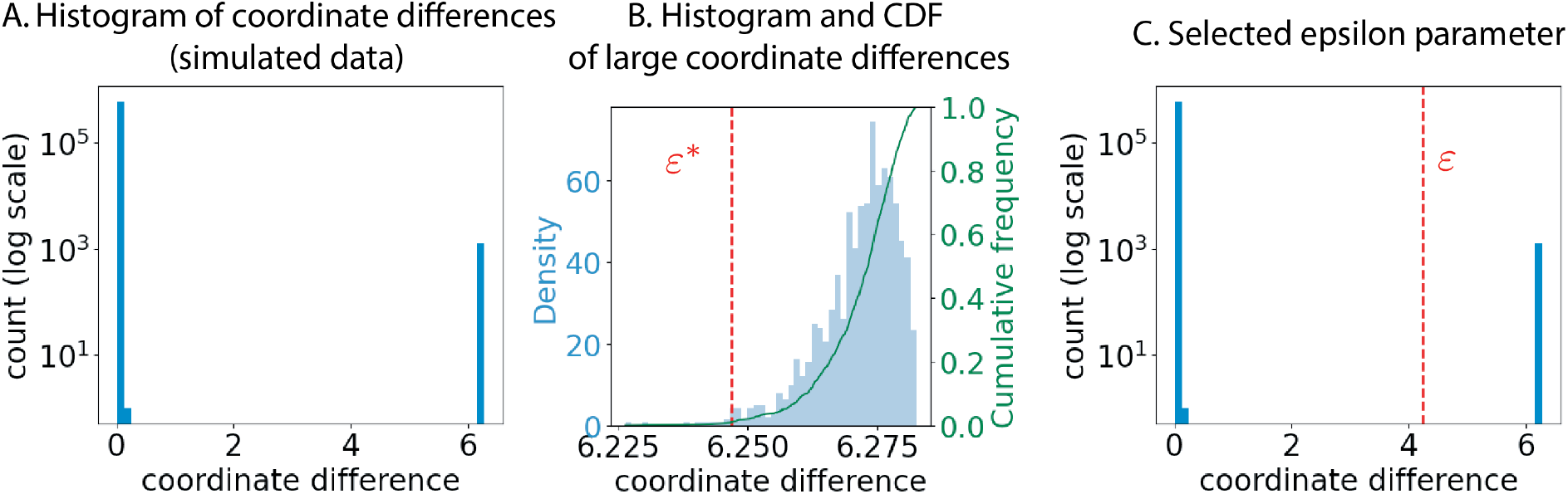
Selection of the proximity parameter *ε* for CAN-simulated data. **A**. Histogram of maximal coordinate differences 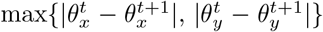 across all consecutive time points, shown on a log scale. The differences concentrate near 0 and 2*π*. **B**. Histogram and empirical cumulative distribution function (green curve) of the coordinate differences restricted to the interval [2, 2*π*). The red dashed line indicates the threshold *ε*^∗^ at which 99% of the maximal coordinate differences exceed *ε*^∗^. **C**. The final parameter is set to *ε* = *ε*^∗^ − 2.

Concretely, we restrict the maximal coordinate difference to [2, 2*π*) (Fig. 9B, SI Fig. 9B, center) and compute their empirical cumulative distribution function. We select *ε*^∗^ to be the value at which *P*(*X > ε*^∗^) = 0.99, ensuring that 99% of the larger coordinate differences lie above *ε*^∗^. We then set the final parameter to *ε* = *ε*^∗^ − 2, which provides additional margin for identifying potential lifts (Fig. 9C, SI Fig. 9B, right). The time points whose toroidal coordinate differences are greater than this *ε* are potential time points for nontrivial lifts. Whether the nontrivial lifts occur, and if so, in what manner, is determined by Equation 3.

#### 4.3.4. Null baselines for assessing reconstruction quality

To assess whether the reconstructed paths are meaningfully similar to the original trajectories, we compare the reconstruction errors against null baselines that quantify the expected error when no true correspondence exists between two paths. We use different null constructions for the simulated and experimental settings.

##### Simulated data

For each environment (0, 1, and 2 holes), we independently simulated pairs of trajectories using the same random walk model and environment parameters described in *Materials and Methods* Section 4.1.1. For each pair, we computed the optimal affine transformation aligning one trajectory to the other and computed the error as described in *Materials and Methods* Section 4.3.2. This process was repeated 100 times to obtain a distribution of null errors. Because the two trajectories in each pair are generated independently, any alignment between them is purely coincidental, and the resulting error distribution represents the expected reconstruction error in the absence of a meaningful relationship between the reconstructed and original paths (Fig. 4E).

##### Experimental data (two-dimensional)

To assess the quality of the reconstruction on local paths, we constructed a null baseline from mismatched segment pairs. Specifically, given the set of all 20-second local path segments, for each pair of path segments, we aligned one trajectory to the other and computed the error as described in *Materials and Methods* Section 4.3.2. There were 63 local path segments, so we obtained a total of 1,953 baseline errors. The distribution of these errors serves as a baseline against which the true (matched) reconstruction errors are compared (Fig. 7K). A reconstruction error significantly lower than this baseline indicates that the reconstructed path preserves the geometry of the specific original segment from which it was derived, rather than reflecting generic properties shared across all segments.

##### Null models for path lifting

To verify that the inductive lifting procedure is necessary for accurate path reconstruction, we compared the proposed method against two null models: a no-lifting model, in which the toroidal coordinates are used directly as the reconstructed path without any tile transitions, and a random-lifting model, in which the coordinates at each time point are lifted to a tile chosen randomly among the preceding lift’s tile and its eight neighbors. Both null models were applied to simulated trajectories in the one-hole environment. See SI Section 2 for details.

## 4.4. Code Availability

Code can be found at the following Github repository https://github.com/irishryoon/GridPathLifting. For simulations, we modified the code shared in https://github.com/erikher/GridCellTorus.

All computations were performed on a single core of an Intel Xeon Gold 6326 (2.90 GHz) node with 512 GB RAM. For a single simulated dataset (2,464 neurons, 600,000 time bins), the full pipeline was completed in approximately 12 minutes (3 minutes for grid cell activity simulation, 9 minutes for the toroidal coordinates computation, 0.5 seconds for path lifting). For the two-dimensional experimental data (111 neurons, 126,728 time bins), the full pipeline was completed in less than a minute. For reference, the same pipeline required approximately 80 minutes for the simulated data and less than a minute for the experimental data on a standard laptop (MacBook Pro, Apple M2, 16 GB RAM, single core).

## 4.5. Data Availability

All experimental data in this manuscript are from publicly available datasets. The one-dimensional movement dataset [44] can be found at https://data.mendeley.com/datasets/rgtk6jygjc/1.

We analyzed the dataset “N2 200203 buildup track.mat,” which contains recordings from mouse N2 navigating a 320 cm virtual build-up track. The dataset provides spatially binned firing rates (2-cm bins, *T* = 160 bins per lap) for *G* = 44 co-modular grid cells identified via clustering on power spectra. We analyzed 441 continuous runs on the track.

The two-dimensional open field movement dataset [20] can be found at https://figshare.com/articles/dataset/Toroidal_topology_of_population_activity_in_grid_cells/16764508.

We analyzed data from rat R, module 1, day 2, open-field session, in which the rat explored a 1.5 m × 1.5 m square arena for 21.1 minutes. Spikes from grid cells in layers II and III of MEC-parasubiculum were recorded. Six grid modules were identified by clustering; we analyzed *G* = 111 co-modular grid cells. Spike trains were preprocessed into continuous firing rate estimates as described in Section 4.2.2. The resulting firing rates were reported over *T* = 126, 728 time bins, where 1 time bin corresponds to 10 ms.

Both datasets have been made public by the authors of the corresponding publications.

## Footnotes

1 A torus is a space that represents the outside surface of a donut. Here, we represent a torus by identifying the left and right edges and by identifying the top and bottom edges of a parallelogram

2 In general, *T* represents the number of time bins. In some cases, as in the one-dimensional experimental data from [44] analyzed in this paper, grid cell activity is presented over spatial bins, in which case *T* represents the number of spatial bins.

3 The entries of the matrix can be binary, with *A*(*i, j*) = 1 indicating that neuron *i* fired at time bin *j* and *A*(*i, j*) = 0 representing that neuron *i* did not fire at time bin *j*, or non-negative real numbers, with *A*(*i, j*) representing the activity level of neuron *i* at time bin *j*. In this work, the entries represent activity values.

4 If two consecutive time points *t* and *t* + 1 are far apart, possibly because the time sampling is too sparse, then the subject could have traveled a large distance between them. Then, the lifted point 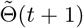 may need to be placed in a tile that is not adjacent to the tile containing 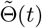. Here, we assume that temporal sampling is dense enough relative to the subject’s movement speed that consecutive population vectors always correspond to the same or adjacent tiles.

## References

[1] Andrea Banino, Caswell Barry, Benigno Uria, Charles Blundell, Timothy Lillicrap, Piotr Mirowski, Alexander Pritzel, Martin J. Chadwick, Thomas Degris, Joseph Modayil, Greg Wayne, Hubert Soyer, Fabio Viola, Brian Zhang, Ross Goroshin, Neil Rabinowitz, Razvan Pascanu, Charlie Beattie, Stig Petersen, Amir Sadik, Ivo Danihelka, Martin Riedmiller, Andreas Fidjeland, Dominik Grewe, Demis Hassabis, and Dharshan Kumaran. “Vector-based navigation using grid-like representations in artificial agents”. In: Nature 557.7705 (2018), pp. 429–433. DOI: 10.1038/s41586-018-0102-6.

[2] Caswell Barry and Daniel Bush. “From A to Z: a potential role for grid cells in spatial navigation”. In: Neural Systems & Circuits 2.1 (May 2012), p. 6. DOI: 10.1186/2042-1001-2-6.

[3] Ulrich Bauer. “Ripser: efficient computation of Vietoris–Rips persistence barcodes”. In: Journal of Applied and Computational Topology 5.3 (2021), pp. 391–423.

[4] G. Bradski. “The OpenCV Library”. In: Dr. Dobb’s Journal of Software Tools (2000).

[5] Yoram Burak and Ila R Fiete. “Accurate path integration in continuous attractor network models of grid cells”. In: PLoS computational biology 5.2 (2009), e1000291.

[6] Daniel Bush, Caswell Barry, Daniel Manson, and Neil Burgess. “Using grid cells for navigation”. In: Neuron 87.3 (2015), pp. 507–520.

[7] Gunnar Carlsson. “Topology and Data”. In: Bulletin of The American Mathematical Society - BULL AMER MATH SOC 46 (Apr. 2009), pp. 255–308. DOI: 10.1090/S0273-0979-09-01249-X.

[8] Rishidev Chaudhuri, Berk Gerçek, Biraj Pandey, Adrien Peyrache, and Ila Fiete. “The intrinsic attractor manifold and population dynamics of a canonical cognitive circuit across waking and sleep”. en. In: Nature Neuroscience 22.9 (Sept. 2019). Number: 9 Publisher: Nature Publishing Group, pp. 1512–1520. ISSN: 1546-1726. DOI: 10.1038/s41593-019-0460-x.

[9] Jonathan J Couey, Aree Witoelar, Sheng-Jia Zhang, Kang Zheng, Jing Ye, Benjamin Dunn, Rafal Czajkowski, May-Britt Moser, Edvard I Moser, Yasser Roudi, et al. “Recurrent inhibitory circuitry as a mechanism for grid formation”. In: Nature neuroscience 16.3 (2013), pp. 318–324.

[10] Carina Curto and Vladimir Itskov. “Cell Groups Reveal Structure of Stimulus Space”. In: PLoS Computational Biology 4.10 (2008), e1000205. DOI: 10.1371/journal.pcbi.1000205.

[11] Yuri Dabaghian. “Grid Cell Percolation”. In: Neural Computation 35.10 (2023), pp. 1609–1626.

[12] Suogui Dang, Yining Wu, Rui Yan, and Huajin Tang. “Why grid cells function as a metric for space”. In: Neural Networks 142 (2021), pp. 128–137.

[13] Herbert Edelsbrunner, John Harer, et al. “Persistent homology-a survey”. In: Contemporary mathematics 453.26 (2008), pp. 257–282.

[14] Edelsbrunner, Letscher, and Zomorodian. “Topological persistence and simplification”. In: Discrete & computational geometry 28 (2002), pp. 511–533.

[15] Uğur M Erdem and Michael Hasselmo. “A goal-directed spatial navigation model using forward trajectory planning based on grid cells”. In: European Journal of Neuroscience 35.6 (2012), pp. 916–931.

[16] Ila R. Fiete, Yoram Burak, and Ted Brookings. “What Grid Cells Convey about Rat Location”. en. In: Journal of Neuroscience 28.27 (July 2008). Publisher: Society for Neuroscience Section: Articles, pp. 6858–6871. ISSN: 0270-6474, 1529-2401. DOI: 10.1523/JNEUROSCI.5684-07.2008.

[17] Martin A. Fischler and Robert C. Bolles. “Random Sample Consensus: A Paradigm for Model Fitting with Applications to Image Analysis and Automated Cartography”. In: Communications of the ACM 24.6 (1981), pp. 381–395. DOI: 10.1145/358669.358692.

[18] Markus Frey, Sander Tanni, Catherine Perrodin, Alice O’Leary, Matthias Nau, Jack Kelly, Andrea Banino, Christian F. Doeller, and Caswell Barry. Deepinsight: a general framework for interpreting wide-band neural activity. en. Pages: 871848 Section: New Results. Dec. 2019. DOI: 10.1101/871848.

[19] Mark C Fuhs and David S Touretzky. “A spin glass model of path integration in rat medial entorhinal cortex”. In: Journal of Neuroscience 26.16 (2006), pp. 4266–4276.

[20] Richard J. Gardner, Erik Hermansen, Marius Pachitariu, Yoram Burak, Nils A. Baas, Benjamin A. Dunn, May-Britt Moser, and Edvard I. Moser. “Toroidal topology of population activity in grid cells”. In: Nature 602.7895 (Feb. 2022). Publisher: Nature Publishing Group, pp. 123–128. ISSN: 1476-4687. DOI: 10.1038/s41586-021-04268-7.

[21] Robert Ghrist. “Barcodes: the persistent topology of data”. In: Bulletin of the American Mathematical Society 45.1 (2008), pp. 61–75.

[22] Torkel Hafting, Marianne Fyhn, Sturla Molden, May-Britt Moser, and Edvard I Moser. “Microstructure of a spatial map in the entorhinal cortex”. In: Nature 436.7052 (2005), pp. 801–806.

[23] D. H. Hubel and T. N. Wiesel. “Receptive fields and functional architecture of monkey striate cortex”. eng. In: The Journal of Physiology 195.1 (Mar. 1968), pp. 215–243. ISSN: 0022-3751. DOI: 10.1113/jphysiol.1968.sp008455.

[24] Jesse A Livezey and Joshua I Glaser. “Deep learning approaches for neural decoding across architectures and recording modalities”. In: Briefings in Bioinformatics 22.2 (Dec. 2020). eprint: https://academic.oup.com/bib/article-pdf/22/2/1577/36654842/bbaa355.pdf, pp. 1577– 1591. ISSN: 1477-4054. DOI: 10.1093/bib/bbaa355.

[25] Cécile Masson and Benoît Girard. “Decoding the Grid Cells for Metric Navigation Using the Residue Numeral System”. In: Advances in Cognitive Neurodynamics (II). Ed. by Rubin Wang and Fanji Gu. Dordrecht: Springer Netherlands, 2011, pp. 459–464. ISBN: 978-90-481-9695-1.

[26] Alexander Mathis, Andreas VM Herz, and Martin Stemmler. “Optimal population codes for space: grid cells outperform place cells”. In: Neural computation 24.9 (2012), pp. 2280–2317.

[27] Bruce L McNaughton, Francesco P Battaglia, Ole Jensen, Edvard I Moser, and May-Britt Moser. “Path integration and the neural basis of the’cognitive map’”. In: Nature Reviews Neuroscience 7.8 (2006), pp. 663–678.

[28] Edward C. Mitchell, Brittany Story, David Boothe, Piotr J. Franaszczuk, and Vasileios Maroulas. “A topological deep learning framework for neural spike decoding”. English. In: Biophysical Journal 123.17 (Sept. 2024). Publisher: Elsevier, pp. 2781–2789. ISSN: 0006-3495, 1542-0086. DOI: 10.1016/j.bpj.2024.01.025.

[29] J. Munkres. Topology. Pearson Modern Classics for Advanced Mathematics Series. Pearson, 2017. ISBN: 9780134689517.

[30] John O’Keefe. “Place units in the hippocampus of the freely moving rat”. In: Experimental Neurology 51.1 (Jan. 1976), pp. 78–109. ISSN: 0014-4886. DOI: 10.1016/0014-4886(76)90055-8.

[31] Jing-Jie Peng, Beate Throm, Maryam Najafian Jazi, Ting-Yun Yen, Hannah Monyer, and Kevin Allen. Grid cells perform path integration in multiple reference frames during self-motion-based navigation. en. Pages: 2023.12.21.572857 Section: New Results. Dec. 2023. DOI: 10.1101/2023.12.21.572857.

[32] Jose A. Perea, Luis Scoccola, and Christopher J. Tralie. “DREiMac: Dimensionality Reduction with Eilenberg-MacLane Coordinates”. In: Journal of Open Source Software 8.91 (2023), p. 5791. DOI: 10.21105/joss.05791.

[33] Erik Rybakken, Nils Baas, and Benjamin Dunn. “Decoding of Neural Data Using Cohomological Feature Extraction”. In: Neural Computation 31.1 (Jan. 2019), pp. 68–93. ISSN: 0899-7667.

[34] Luis Scoccola, Hitesh Gakhar, Johnathan Bush, Nikolas Schonsheck, Tatum Rask, Ling Zhou, and Jose A. Perea. “Toroidal Coordinates: Decorrelating Circular Coordinates with Lattice Reduction”. In: LIPIcs, Volume 258, SoCG 2023 258 (2023). In collab. with Erin W. Chambers and Joachim Gudmundsson. Artwork Size: 20 pages, 6894838 bytes ISBN: 9783959772730 Medium: application/pdf Publisher: Schloss Dagstuhl – Leibniz-Zentrum für Informatik, 57:1–57:20. ISSN: 1868-8969. DOI: 10.4230/LIPICS.SOCG.2023.57.

[35] Vin de Silva, Dmitriy Morozov, and Mikael Vejdemo-Johansson. “Persistent Cohomology and Circular Coordinates”. en. In: Discrete & Computational Geometry 45.4 (June 2011), pp. 737–759. ISSN: 1432-0444. DOI: 10.1007/s00454-011-9344-x.

[36] Arturs Simkuns, Rodions Saltanovs, Maksims Ivanovs, and Roberts Kadikis. “Deep Learning-Emerged Grid Cells-Based Bio-Inspired Navigation in Robotics”. In: Sensors 25.5 (2025), p. 1576. DOI: 10.3390/s25051576.

[37] Trygve Solstad, Edvard I Moser, and Gaute T Einevoll. “From grid cells to place cells: a mathematical model”. In: Hippocampus 16.12 (2006), pp. 1026–1031.

[38] Martin Stemmler, Alexander Mathis, and Andreas VM Herz. “Connecting multiple spatial scales to decode the population activity of grid cells”. In: Science Advances 1.11 (2015), e1500816.

[39] Hong Sun and Tian-Ren Yao. “A neural-like network approach to residue-to-decimal conversion”. In: Proceedings of 1994 IEEE International Conference on Neural Networks (ICNN’94). Vol. 6. June 1994, 3883–3887 vol.6. DOI: 10.1109/ICNN.1994.374831.

[40] Ardi Tampuu, Tambet Matiisen, H. Freyja Ólafsdóttir, Caswell Barry, and Raul Vicente. “Efficient neural decoding of self-location with a deep recurrent network”. en. In: PLOS Computational Biology 15.2 (Feb. 2019). Publisher: Public Library of Science, e1006822. ISSN: 1553-7358. DOI: 10.1371/journal.pcbi.1006822.

[41] Jeffrey S Taube, Robert U Muller, and James B Ranck. “Head-direction cells recorded from the postsubiculum in freely moving rats. I. Description and quantitative analysis”. In: Journal of Neuroscience 10.2 (1990), pp. 420–435.

[42] Christopher Tralie, Nathaniel Saul, and Rann Bar-On. “Ripser. py: A lean persistent homology library for python”. In: Journal of Open Source Software 3.29 (2018), p. 925.

[43] Vincent Villette, Arnaud Malvache, Thomas Tressard, Nathalie Dupuy, and Rosa Cossart. “Internally Recurring Hippocampal Sequences as a Population Template of Spatiotemporal Information”. eng. In: Neuron 88.2 (Oct. 2015), pp. 357–366. ISSN: 1097-4199. DOI: 10.1016/j.neuron.2015.09.052.

[44] John H Wen, Ben Sorscher, Emily A Aery Jones, Surya Ganguli, and Lisa M Giocomo. “One-shot entorhinal maps enable flexible navigation in novel environments”. In: Nature 635.8040 (2024), pp. 943–950.

[45] Zishen Xu, Wei Wu, Shawn S. Winter, Max L. Mehlman, William N. Butler, Christine M. Simmons, Ryan E. Harvey, Laura E. Berkowitz, Yang Chen, Jeffrey S. Taube, Aaron A. Wilber, and Benjamin J. Clark. “A Comparison of Neural Decoding Methods and Population Coding Across Thalamo-Cortical Head Direction Cells”. English. In: Frontiers in Neural Circuits 13 (Dec. 2019). Publisher: Frontiers. ISSN: 1662-5110. DOI: 10.3389/fncir.2019.00075.

[46] Yuansheng Zhou, Brian H. Smith, and Tatyana O. Sharpee. “Hyperbolic geometry of the olfactory space”. In: Science Advances 4.8 (Aug. 2018). Publisher: American Association for the Advancement of Science, eaaq1458. DOI: 10.1126/sciadv.aaq1458.

[47] Qiang Zou, Ming Cong, Dong Liu, and Yu Du. “A neurobiologically inspired mapping and navigating framework for mobile robots”. In: Neurocomputing 460 (Oct. 2021), pp. 181–194. DOI: 10.1016/j.neucom.2021.07.025.

