## Supplementary Information for "Topological decoding of grid cell activity via path lifting to covering spaces"

### Contents

1. Mathematical Preliminaries 1
1.1. Lifting paths on torus to paths in $\mathbb{R}^2$ 1
1.2. Persistent homology 2
2. Comparison against null models of lifting 5
3. Robustness against noise in neural activity 6
3.1. Robustness against spontaneous activity 6
3.2. Robustness against neural activity suppression 9
3.3. Robustness against time shifts of neural activity 11
4. Impact of various factors on global and local reconstruction errors 13
4.1. Impact of proximity parameter epsilon 13
4.2. Impact of number of time points 14
4.3. Error accumulation for long paths and impact of experiment duration 17
4.4. Impact of number of neurons 20
4.5. Impact of metric 20
4.6. Impact of noise in toroidal coordinates 21
4.7. Impact of smoothing reconstructed paths from experimental data 22
5. Supplementary Figures 24
References 26

### 1. Mathematical Preliminaries

1.1. **Lifting paths on torus to paths in $\mathbb{R}^2$.** Given a topological space $X$, a path in $X$ is a continuous map $f : I \to X$, where $I = [0, 1]$ denotes the unit interval. If one considers $t \in [0, 1]$ as representing time, then $f(t)$ specifies the location of an object in $X$ at time $t$.

In this work, we are concerned with lifting paths on a torus to paths in $\mathbb{R}^2$. Recall that $S^1$ represents a circle, and that $S^1 \times S^1$ represents a torus. Let $f : I \to S^1 \times S^1$ be a path on the torus, and let $p : \mathbb{R}^2 \to S^1 \times S^1$ be a covering of a torus. For example, $p : \mathbb{R}^2 \to S^1 \times S^1$ defined by

$$p(x, y) = (\,(\cos 2\pi x, \sin 2\pi x),\ (\cos 2\pi y, \sin 2\pi y)\,) \tag{1}$$

is a valid covering of a torus. Note that $(\cos 2\pi x, \sin 2\pi x)$ and $(\cos 2\pi y, \sin 2\pi y)$ each specify points in $S^1$.

The following lemma states that any path on a torus can be lifted to a path in $\mathbb{R}^2$.

**Lemma 1.** *(Lemma 54.2 [6], modified) The path $f : I \to S^1 \times S^1$ can be lifted to a path $\tilde{f} : I \to \mathbb{R}^2$ such that the following diagram commutes.*

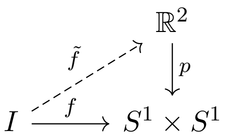


*Furthermore, given a $b_0 = f(0) \in S^1 \times S^1$ and $e_0 \in p^{-1}(b_0)$, the lifted path $\tilde{f}$ with $\tilde{f}(0) = e_0$ is unique.*

A constructive proof can be found in [6]. Here, we illustrate the construction of $\tilde{f}$ in a simple example. Let $p : \mathbb{R}^2 \to S^1 \times S^1$ the covering map from Equation 1. We construct $\tilde{f}$ in pieces.

Let $U_1, \dots, U_4$ be open sets covering the torus (Fig. 1A). Note that the for any $U \in \{U_1, \dots U_4\}$, the preimage $p^{-1}(U)$ consists of infinitely-many homeomorphic copies of $U$ in $\mathbb{R}^2$ (SI Fig. 1B). Each copy of $U$ in $p^{-1}(U)$ is called a *slice*.

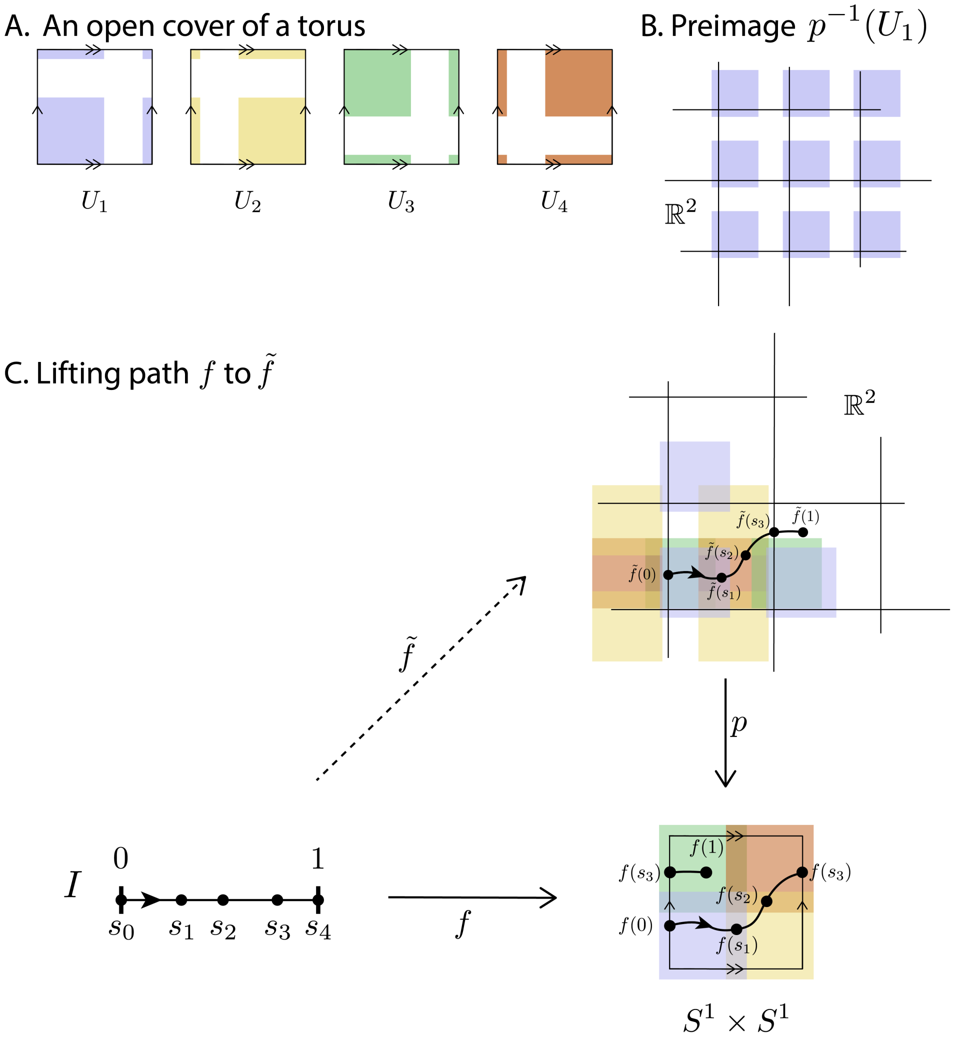


**SI Figure 1.** Lifting a path $f$ on torus to a path $\tilde{f}$ in $\mathbb{R}^2$. **A.** The open cover $\{U_1, \dots, U_4\}$ of the torus. **B.** The preimage $p^{-1}(U_1)$ consists of homeomorphic copies of $U_1$ in $\mathbb{R}^2$. **C.** The lifted path $\tilde{f}$ is constructed piece-by-piece.

We then partition $I = [0, 1]$ into segments $0 = s_0 < s_1 < \dots < s_4 = 1$ so that $f$ maps each segment $[s_i, s_{i+1}]$ into one of $U_1, \dots, U_4$ (Figure 1C). Without loss of generality, assume $f(0)$ lies in $U_1$. We choose some slice, let's say $V_1$, of $U_1$. We let $\tilde{f}(0)$ be the unique point in $V_1^1$ that maps to $f(0)$ via $p$. We then define $\tilde{f}$ on $[0, s_1]$ to be the unique path in $V_1 \subset \mathbb{R}^2$ that maps to $f|_{[0,s_1]}$ via $p$.

To define $\tilde{f}$ on $[s_1, s_2]$, note that $f(s_1)$ lives in both $U_1$ and $U_2$. Among the slices $p^{-1}(U_2)$ of $U_2$, there exists a unique slice, say $V_2$ of $U_2$ where $\tilde{f}(s_1) \in V_2$. We then define $\tilde{f}$ on $[s_1, s_2]$ to be the unique path in $V_2 \subset \mathbb{R}^2$ that maps to $f|_{[s_1,s_2]}$ via $p$.

We continue this procedure until we define $\tilde{f}$ on the entire interval $[0, 1]$. By construction, $p \circ \tilde{f} = f$.

1.2. **Persistent homology.** We provide a brief description of simplicial homology (with field coefficients) and persistent homology.

1.2.1. *Simplicial Complexes and Simplicial Homology.* An *(abstract) simplicial complex* $K = (V, F)$ is a combinatorial structure built from a set of vertices $V$. It consists of simplices, where a simplex is an unordered subset of $V$. A collection of $n+1$ elements in $V$, for example, $(v_0, \dots, v_n)$, is called an $n$-simplex. Concretely, a single vertex is a 0-simplex, the collection $(v_0, v_1)$ is a 1-simplex, and $(v_0, v_1, v_2)$ is a 2-simplex. The collection of simplices $F$ must satisfy the following: given a simplex $\sigma \in F$, all of its non-empty subsets must also be in $F$.

In this paper, the homology of a simplicial complex is computed with field coefficients $\mathbb{F}$. Therefore, all homology computations are done in the context of vector spaces and linear maps.

To study the topology of $K$, we work with *chains*, which are formal linear combinations of simplices. More precisely, if $F_n$ denotes the number of $n$-simplices in a complex $K$, then the vector space of $n$-chains is

$$C_n(K) = \Big\{ \sum_{i=1}^{F_n} c_i \sigma_i^n \;\Big|\; c_i \in \mathbb{F},\ \sigma_i^n \in K \Big\}.$$

The *boundary homomorphism* $\partial_n : C_n(K) \to C_{n-1}(K)$ is constructed as follows. For an $n$-simplex $\sigma = (v_0, \dots, v_n)$, the boundary is defined as

$$\partial_n(\sigma) = \sum_{i=0}^{n} (-1)^i (v_0, \dots, \hat{v}_i, \dots, v_n),$$

where the notation $\hat{v}_i$ means that the vertex $v_i$ has been omitted. This map is then extended linearly to all of $C_n(K)$. We then obtain the following

$$\cdots \to C_{n+1}(K) \xrightarrow{\partial_{n+1}} C_n(K) \xrightarrow{\partial_n} C_{n-1}(K) \to \cdots \to C_0(K) \to 0,$$

which is a sequence of vector spaces and linear maps. The boundary homomorphisms satisfy the key property that $\partial_n \circ \partial_{n+1} = 0$ for all $n$.

This property ensures that $\operatorname{im} \partial_{n+1} \subseteq \ker \partial_n$. Elements in $\ker \partial_n$ are called *cycles*, representing potential $n$-dimensional holes, while those in $\operatorname{im} \partial_{n+1}$ are called *boundaries*, representing cycles that are filled in by higher-dimensional simplices. The true $n$-dimensional holes are captured by the *homology group*

$$H_n(K) = \ker \partial_n \big/ \operatorname{im} \partial_{n+1}.$$

Each homology group is a vector space, and its dimension records the number of independent $n$-dimensional holes. These invariants provide a compact algebraic summary of the underlying topological structure of the simplicial complex.

1.2.2. *Persistent homology.* We present a short overview of persistent homology. Suppose we have a population $P = \{p_1, \dots, p_n\}$ of interest that we wish to analyze. We assume that the pairwise dissimilarities between any pair $p_i$ and $p_j$ is known. A convenient way to encode the system is through a *simplicial complex* that has $P$ as its vertex set. One way of obtaining this goal is to fix a threshold $\varepsilon > 0$ and construct the simplicial complex $X_P^{\varepsilon} = (P, F_{\varepsilon})$, where the vertices are given by $P$ and a subset $\sigma \subseteq P$ belongs to $F_{\varepsilon}$ whenever all its members are at most $\varepsilon$ apart. The first homology group $H_1(X_P^{\varepsilon})$ then records the 1-dimensional cycles in this complex, with its dimension giving the number of independent loops present.

Choosing a single threshold $\varepsilon$ can be arbitrary, however. Persistent homology addresses this by examining how the homology evolves as $\varepsilon$ varies. Given a sequence of thresholds $\{\varepsilon_1 < \varepsilon_2 < \cdots < \varepsilon_N\}$, one obtains a filtration of simplicial complexes

$$X_P^{\varepsilon_1} \subseteq X_P^{\varepsilon_2} \subseteq \cdots \subseteq X_P^{\varepsilon_N}.$$

Applying homology to this nested sequence yields a diagram of vector spaces and linear maps,

$$H_1(X_P^{\bullet}) :\ H_1(X_P^{\varepsilon_1}) \to H_1(X_P^{\varepsilon_2}) \to \cdots \to H_1(X_P^{\varepsilon_N}), \tag{2}$$

where each map arises from the inclusion of one simplicial complex into another, carrying cycles forward across scales. Persistent homology tracks when a homological feature (such as a cycle) first appears and when it disappears within this filtration.

The lifespan of a feature is summarized by its *birth* parameter $b$ and *death* parameter $d$. Plotting the collection of points $(b, d)$ gives the *persistence diagram*, a compact summary of the multi-scale topological structure. For a comprehensive treatment, see [1, 2, 3, 5]

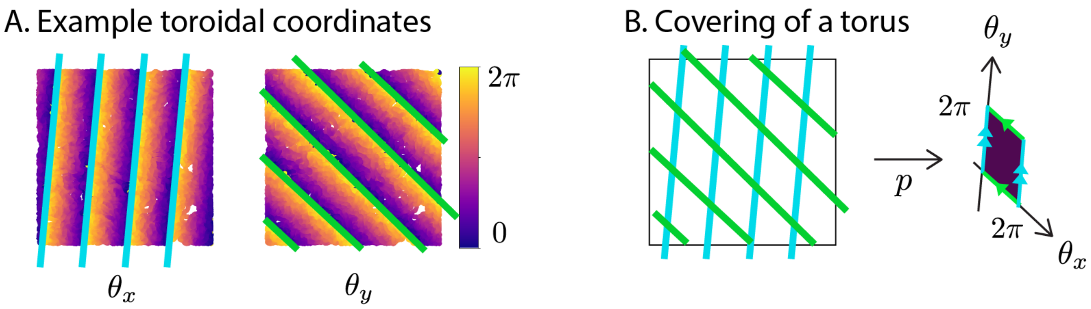


**SI Figure 2.** The toroidal coordinates define a tiling of $\mathbb{R}^2$ via parallelograms. **A**. An example visualization of the toroidal coordinates $(\theta_x, \theta_y)$. Here, given a time point $t$, let $(x, y)$ denote the location of the mouse at time $t$. Let $(\theta_x^t, \theta_y^t)$ be the toroidal coordinate assigned to population vector $P(t)$. The toroidal coordinates are visualized by a scatter plot in which a dot is placed at $(x, y)$ whose color value represents $\theta_x^t$ (left) and $\theta_y^t$ (right). **B**. The toroidal coordinates define a tiling of $\mathbb{R}^2$ via parallelograms. The map $p$ takes each parallelogram to one copy of the grid cell torus $S^1 \times S^1$.

### 2. Comparison against null models of lifting

To demonstrate that the proposed path-lifting procedure is essential to a faithful path reconstruction, we compared the paths reconstructed from our method against two null models: reconstruction without any lifting, and reconstruction with random lifting.

In the first null model (no lifting), the sequence of toroidal coordinates $\{\Theta(t)\}$ is treated directly as the reconstructed path without any lifting. That is, $\tilde{\Theta}(t) = \Theta(t)$, and all lifted coordinates remain in a single tile (see Fig. 2 in main text). In the second null model (random lifting), at each time point $t+1$, the lifted coordinates $\Theta(t+1)$ are assigned to tile randomly chosen from the current tile and its eight neighbors.

We applied all three approaches — the proposed method, no lifting, and random lifting — to simulated trajectories in the one-hole environment (SI Fig. 3). As shown in SI Fig. 3A, only the proposed path-lifting algorithm recovers the hole in the environment; the two null models fail to reproduce the underlying topology of the movement path.

We repeated this experiment 5 times over independent simulations and report the global and local reconstruction errors in SI Fig. 3B. For the local reconstruction error, we take paths of length 10,000 time bins. Consistent with panel A, both null models yield reconstructions that deviate substantially from the true movement path, producing large global and local reconstruction errors.

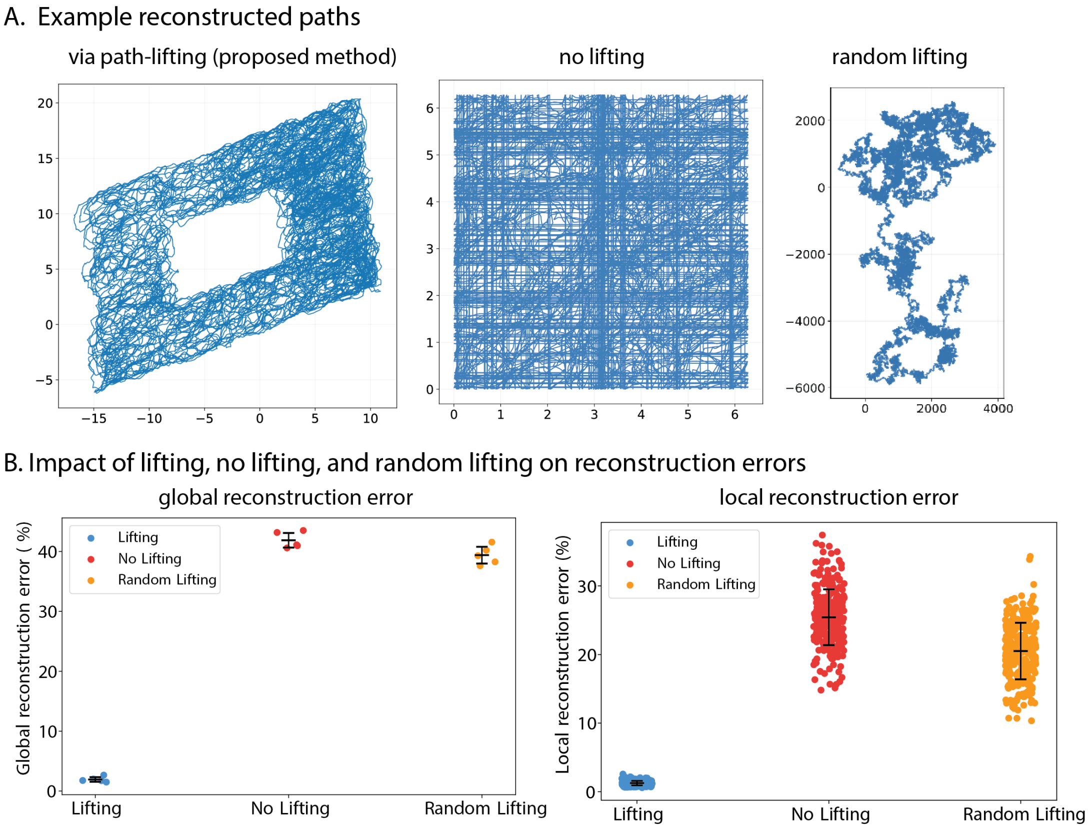


**SI Figure 3.** Comparison of proposed method against two null models (no lifting, random lifting) in the CAN-simulated data. **A.** Example paths reconstructed via the proposed method and two null methods from a simulated trajectory in the one-hole world. **B**. Global (left) and local (right) reconstruction errors across 5 independent simulations. The proposed path-lifting algorithm achieves substantially lower error than both the no-lifting and random-lifting null models. Points show errors from individual simulations; markers and error bars indicate the mean ± 1 standard deviation (for global reconstruction error, mean and standard deviation were computed across 5 independent simulations; for local reconstruction error, they are computed across the 295 local paths, 59 local paths per simulation).

### 3. Robustness against noise in neural activity

In this section, we analyze the robustness of the path reconstruction algorithm against noise in neural activity. We analyze three types of noise: spontaneous activity (SI Section 3.1), neural activity suppression (SI Section 3.2), and time shifts in neural activity (SI Section 3.3) in the CAN-simulated dataset. All reconstruction errors reported in this section are global reconstruction errors.

3.1. **Robustness against spontaneous activity.** We report the global reconstruction errors under varying levels of additional spontaneous activity of simulated grid cells. In the main text, we examine the robustness of the method under addition of one-dimensional Gaussian functions with peak height $h = 0.4$ (main text, Table 1). Here, we examine the reconstruction errors while varying the peak heights $h \in \{0.08, 0.2, 0.3, 0.4\}$. SI Table 1 summarizes the result. See SI Figure 5 for example activity traces after the addition of Gaussian noise of height $h = 0.4$. Note that in Table 1 of main text, we report the mean reconstruction error over 10 independent trials. Here, we report the error from a single trial.

**SI Table 1.** Global reconstruction errors (%) under added Gaussian noise with varying peak heights $h$, proportions $p$ of affected time points, and standard deviations $\sigma$ of the added Gaussian noise. N/A indicates conditions where toroidal coordinates could not be computed.

| | | **Std. Dev.** ($\sigma$) | | | | | | **Std. Dev.** ($\sigma$) | | | |
|---|---|---|---|---|---|---|---|---|---|---|---|
| $h$ | $p$ | 1 | 10 | 50 | 100 | $h$ | $p$ | 1 | 10 | 50 | 100 |
| | 0.1% | 1.96 | 1.68 | 1.78 | 1.78 | | 0.1% | 1.66 | 1.68 | 1.71 | 2.17 |
| | 0.5% | 1.65 | 1.62 | 1.83 | 1.87 | | 0.5% | 1.62 | 1.73 | 25.79 | 45.80 |
| | 1% | 1.64 | 1.71 | 1.87 | 2.11 | | 1% | 1.66 | 1.88 | 71.48 | 41.49 |
| | 5% | 1.62 | 1.84 | 43.69 | N/A | | 5% | 1.71 | 60.36 | N/A | N/A |
| 0.08 | 10% | 1.74 | 2.18 | N/A | N/A | 0.3 | 10% | 16.25 | 40.72 | N/A | N/A |
| | 0.1% | 1.59 | 1.61 | 1.65 | 1.72 | | 0.1% | 1.63 | 1.62 | 1.79 | 11.45 |
| | 0.5% | 1.61 | 1.78 | 2.05 | 32.71 | | 0.5% | 1.63 | 1.82 | 43.88 | 40.82 |
| | 1% | 1.94 | 1.94 | 31.46 | 40.39 | | 1% | 1.86 | 2.06 | 39.47 | 43.11 |
| | 5% | 1.77 | 22.45 | N/A | N/A | | 5% | 7.37 | 58.59 | N/A | N/A |
| 0.2 | 10% | 1.70 | 63.32 | 37.03 | N/A | 0.4 | 10% | 55.23 | 38.31 | N/A | N/A |

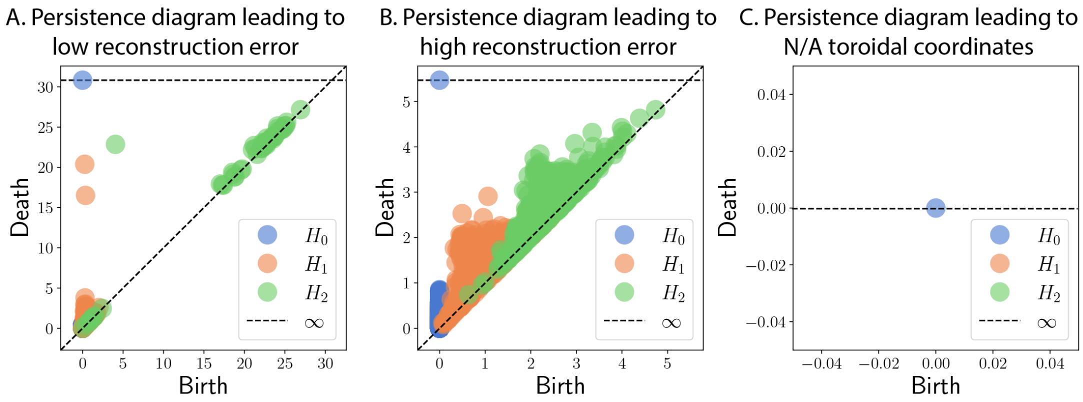


**SI Figure 4.** Example persistence diagrams that lead to low, high, and N/A reconstruction errors in the presence of spontaneous firings with peak height $h = 0.4$. **A**. Persistence diagram of the population vectors for $p = 5\%$ and $\sigma = 1$ has a clear toroidal structure, which leads to a low reconstruction error. **B**. Persistence diagram of the population vectors for $p = 5\%$ and $\sigma = 10$ does not have a clear toroidal structure, which leads to a high reconstruction error. **C**. Persistence diagram of the population vectors for $p = 5\%$ and $\sigma = 100$ has no non-trivial topological features. In such cases, the toroidal coordinates cannot be computed.

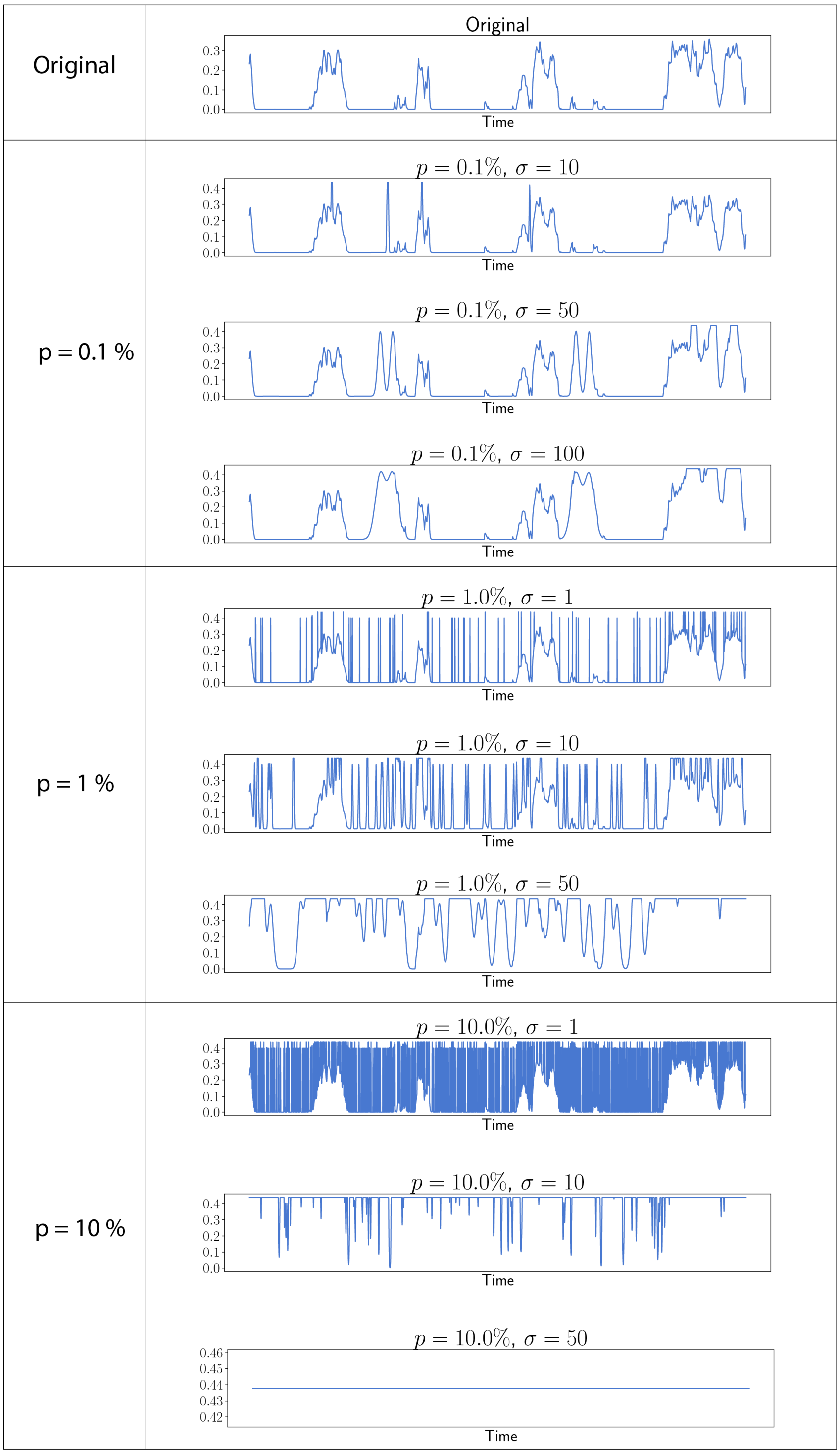


**SI Figure 5.** Simulated grid cell activities with added noise of different parameters of $p$ and $\sigma$. Peak height is fixed at $h = 0.4$.

3.2. **Robustness against neural activity suppression.** To test the robustness of the method against signal dropout or inhibitory transients in neural activity traces, we tested the path reconstruction under varying levels of suppression of simulated grid cell activity. From the CAN-simulated activity, we corrupt each neuron's activity trace as follows. A fraction $p$ (the portion parameter) of time points is selected uniformly at random without replacement. At each selected time point $t_i$, a Gaussian pulse

$$g_i(t) = h \exp\left(-\frac{(t-t_i)^2}{2\sigma^2}\right)$$

is subtracted from the activity trace, where $\sigma$ is the standard deviation controlling the temporal width of the dropout and $h$ is the peak height. The resulting trace is then clipped to $[0, r_{\max}]$, where $r_{\max}$ is the maximum magnitude of the original activity trace, so that no values become negative or exceed the original dynamic range. For varying $h, p$, and $\sigma$ values, we applied this suppression independently to each neuron's activity trace and performed path reconstruction. The results are summarized in SI Table 2. Note that for $h = 0.4$, $p = 10\%$, and $\sigma = 10$, the reconstructed, transformed path occupied a space much larger than a square of size $100 \times 100$. The normalization factor of the reconstruction error (main text, Section 4.3.2) is $S = 100$. Because the transformed path occupies a window larger than $100 \times 100$, dividing by $S = 100$ results in a reconstruction error that is larger than 100%.

See SI Fig. 6 for visualizations of simulated neural activity traces and the modified traces that lead to high and low reconstruction errors.

**SI Table 2.** Reconstruction errors (%) under neural activity suppression for varying peak heights $h$, proportions $p$ of affected time points, and standard deviations $\sigma$ of the subtracted Gaussian pulses. N/A indicates conditions where toroidal coordinates could not be computed.

| | | **Std. Dev.** ($\sigma$) | | | |
|---|---|---|---|---|---|
| $h$ | $p$ | 1 | 10 | 50 | 100 |
| | 0.1% | 1.69 | 1.70 | 1.73 | 1.71 |
| | 0.5% | 1.73 | 1.87 | 1.71 | 1.68 |
| | 1% | 1.65 | 1.66 | 1.67 | 1.73 |
| | 5% | 1.65 | 1.63 | 44.01 | 40.24 |
| 0.08 | 10% | 1.70 | 1.68 | N/A | N/A |
| | 0.1% | 1.68 | 1.67 | 1.66 | 1.72 |
| | 0.5% | 1.67 | 1.59 | 1.90 | 1.82 |
| | 1% | 1.64 | 1.59 | 1.95 | 2.08 |
| | 5% | 1.66 | 1.75 | 43.18 | N/A |
| 0.2 | 10% | 1.59 | 2.08 | N/A | N/A |

| | | **Std. Dev.** ($\sigma$) | | | |
|---|---|---|---|---|---|
| $h$ | $p$ | 1 | 10 | 50 | 100 |
| | 0.1% | 1.63 | 1.60 | 1.68 | 1.64 |
| | 0.5% | 1.75 | 1.64 | 1.77 | 24.09 |
| | 1% | 1.60 | 1.66 | 1.91 | 50.40 |
| | 5% | 1.60 | 1.83 | 66.17 | N/A |
| 0.3 | 10% | 1.62 | 76.02 | N/A | N/A |
| | 0.1% | 1.67 | 1.60 | 1.70 | 1.79 |
| | 0.5% | 1.60 | 1.61 | 1.82 | 2.01 |
| | 1% | 1.64 | 1.66 | 24.39 | 35.65 |
| | 5% | 1.63 | 2.02 | N/A | N/A |
| 0.4 | 10% | 1.67 | 188.44 | N/A | N/A |

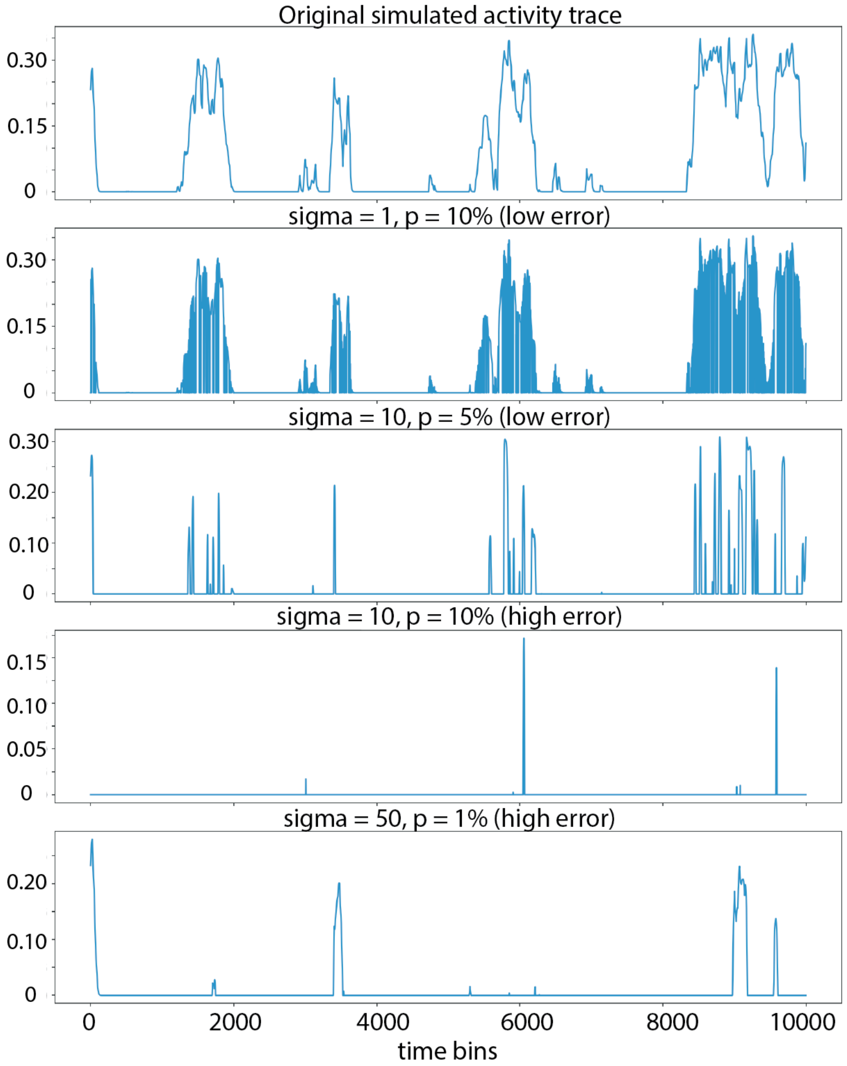


**SI Figure 6.** Original simulated neural activity trace and its modification after neural activity suppression that leads to low and high reconstruction error.

3.3. **Robustness against time shifts of neural activity.** The decoding pipeline assumes that the grid cell activity matrix is temporally aligned, i.e., column $t$ of the activity matrix corresponds to time step $t$ for all neurons simultaneously. To test robustness to violations of this assumption (for instance, arising from clock drift or imprecise spike-sorting alignment across tetrodes), we applied independent random circular shifts to each simulated neuron's activity trace before path reconstruction. Specifically, given a maximum shift amount $d_{\max}$, for each neuron $i$, a temporal shift amount $\delta_i$ was drawn uniformly at random from $[-d_{\max}, d_{\max}]$ (integer-valued), and the trace was cyclically shifted by $\delta_i$ time steps via `np.roll`. Note that the shift amounts are in time bin units, out of a total of 600,000 time bins. See SI Fig. 8 for visualizations of two example neurons and their activity shifted by various time steps.

We considered various maximum shift parameters $d_{\max} \in \{0, 10, 20, 50, 100, 200, 500, 1000\}$. For each shift amount $d_{\max}$, we shifted each neuron's activity trace as described above and performed path reconstruction on the shifted activities. We repeated this experiment on 5 independent simulations.

Both the global and local reconstruction error remained low for maximum shift distance of up to $d_{max} = 100$ time bins, indicating that the pipeline tolerates moderate asynchrony across neurons without appreciable degradation (SI Fig. 7). Performance began to decline noticeably at $d_{max} = 500$. These results demonstrate that the method is robust to small-to-moderate temporal misalignment, while sufficiently large desynchronization disrupts the toroidal structure for path reconstruction.

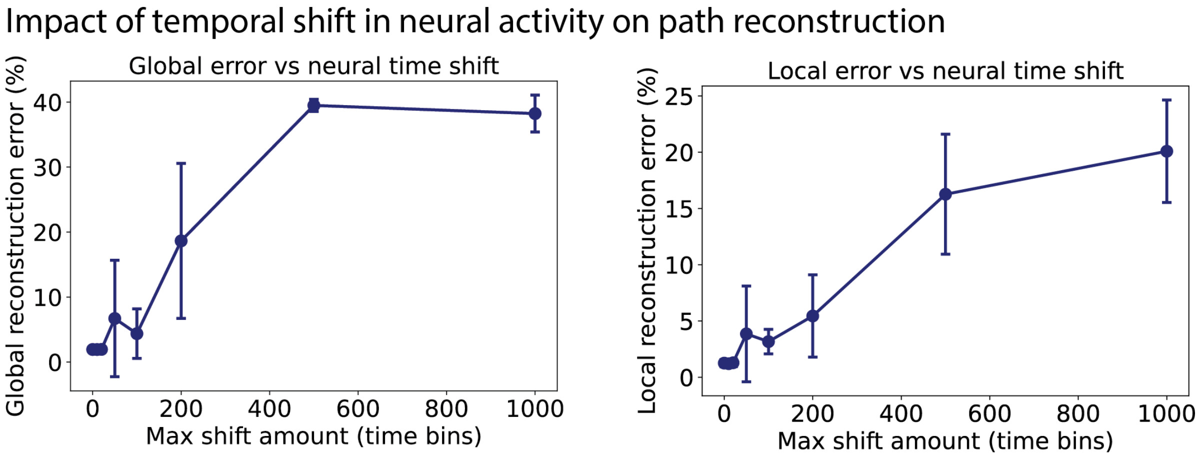


**SI Figure 7.** Effect of temporal shift in neural activity on path reconstruction in simulated data. Each neuron's activity trace was independently circularly shifted by an integer sampled uniformly from $[-d_{\max}, d_{\max}]$, and the shifted population activity was used to perform path reconstruction. Results are shown for 5 independent repeats in the 1-hole simulated environment. Error bars denote standard deviation across repeats. (Left) Global reconstruction error as a function of maximum shift magnitude $d_{\max}$. At $d_{\max} = 500$, the reconstruction error approaches that of the null models. (Right) Local reconstruction error as a function of $d_{\max}$. Each local path was 10,000 time bin long.

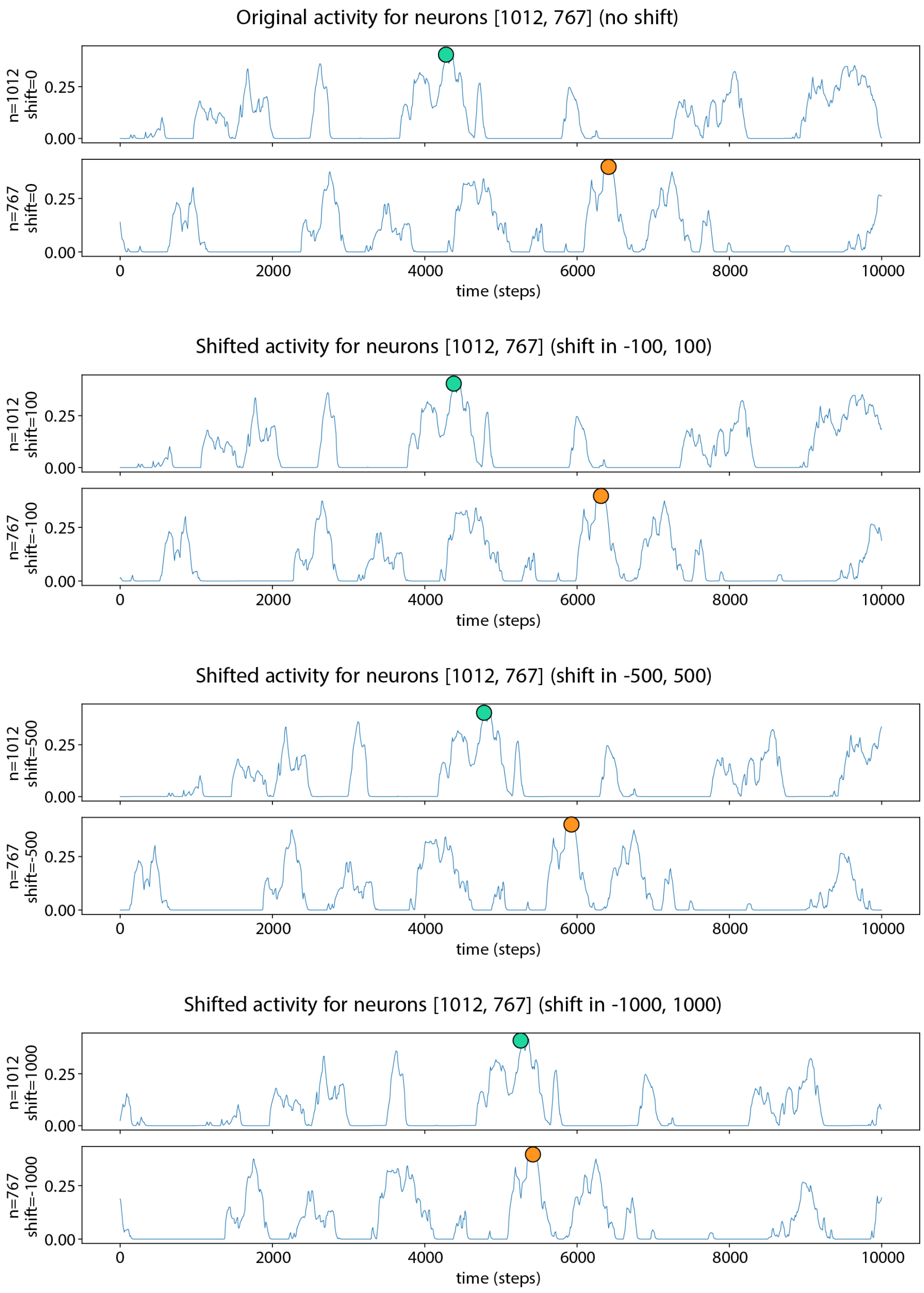


**SI Figure 8.** Example activity of simulated grid cells (neurons 1012 and 767) under deterministic temporal shifts. For each shift magnitude $d_{max} \in \{100, 500, 1000\}$, neuron 1012 is shifted by $+d_{max}$ and neuron 767 by $-d_{max}$, and the first 10,000 time points are shown to illustrate how increasing shift size alters the alignment of the two neural activities. For ease of comparison, the maximal neural activities of each neuron is marked with teal and orange circles.

### 4. Impact of various factors on global and local reconstruction errors

In this section, we analyze the various factors that can potentially impact global and local reconstruction errors. The factors analyzed are proximity parameter epsilon (SI Section 4.1), number of time points (SI Section 4.2), experiment duration (SI Section 4.3), number of neurons (SI Section 4.4), metric (SI Section 4.5), noise in toroidal coordinates (SI Section 4.6), and smoothing of reconstructed paths (SI Section 4.7).

Throughout this section, all analysis was performed on five independent simulations on one-hole environment. When presenting the global reconstruction errors, the points and error bars indicate the mean and standard deviation across the five simulations. For the local reconstruction errors, all local paths had length 10,000 time bins (out of 599,999 total time bins). The mean and standard deviation are computed across all 295 local paths (59 local paths per simulation).

Whenever applicable, we also performed an analogous analysis on the two-dimensional experimental data [4] (111 grid cells from rat R, module 1, day 2, open-field session). The local paths of the two-dimensional experimental data consisted of 20-second intervals. In all figures reporting the local reconstruction errors from the two-dimensional experimental dataset, the points and error bars indicate the mean and standard deviation across all local paths.

4.1. **Impact of proximity parameter epsilon.** In the path reconstruction algorithm, whether two consecutive toroidal coordinates should be tested for potential lifts is determined by a proximity parameter $\varepsilon$ (Equation 2, main text). We analyzed the impact of varying this parameter on the simulated dataset (SI Fig. 9A, left) and the two-dimensional experimental dataset (SI Fig. 9A, center, right). Smaller epsilon parameters lead to lower reconstruction errors, and the errors are stable for a wide range of epsilon parameters.

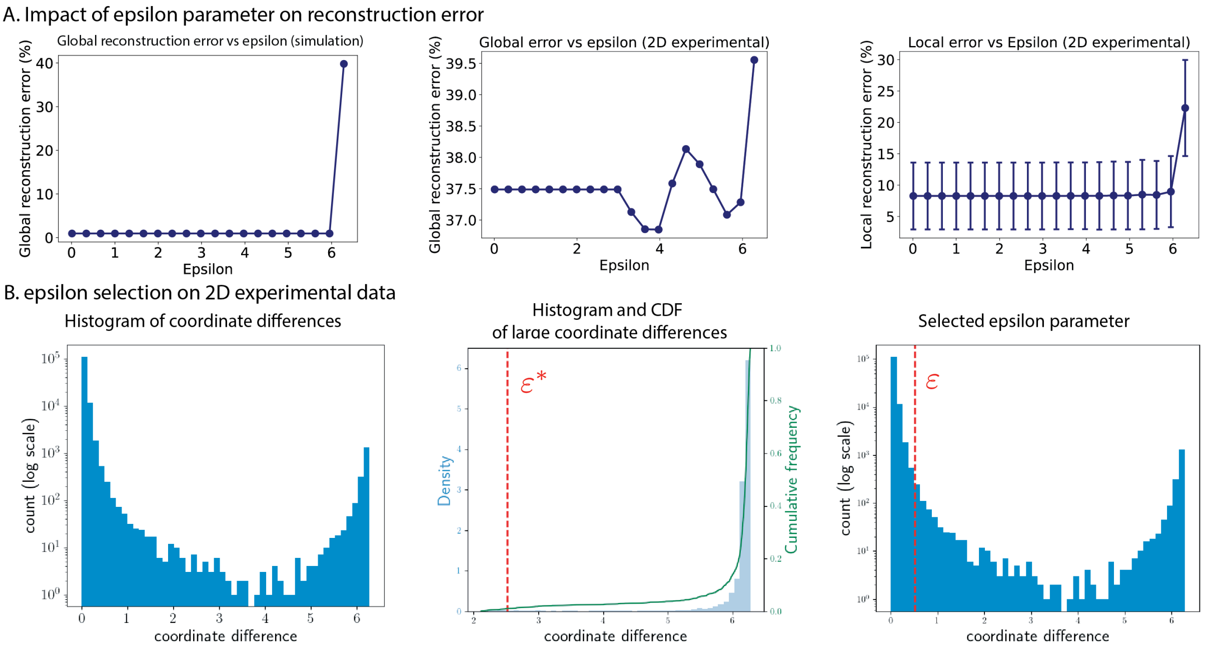


**SI Figure 9.** Impact of proximity parameter $\varepsilon$ on reconstruction error and selection of $\varepsilon$. **A.** Analysis of impact of epsilon parameter on the global reconstruction errors on the CAN-simulated dataset (left) and in two-dimensional experimental data (center). Impact of epsilon parameter on the local reconstruction error in two-dimensional experimental data (right). **B.** $\varepsilon$ selection for two-dimensional experimental data. (Left) A histogram of the maximal coordinate differences. (Center) A zoomed in view of the coordinate differences restricted to the range $[2, 2\pi)$. The green curve shows the cumulative sum. The red dotted line indicates the parameter $\varepsilon^*$ at which 99% of the maximal coordinate differences exceed $\varepsilon^*$. (Right) As observed in panel A, smaller epsilon parameters generally lead to better reconstruction. We choose $\varepsilon = \varepsilon^* - 2$.

4.2. **Impact of number of time points.** A faithful path reconstruction requires that the grid cell population activity is observed at sufficient number of time points. Having insufficient number of time points can lead to poor path reconstructions as illustrated in SI Fig. 10.

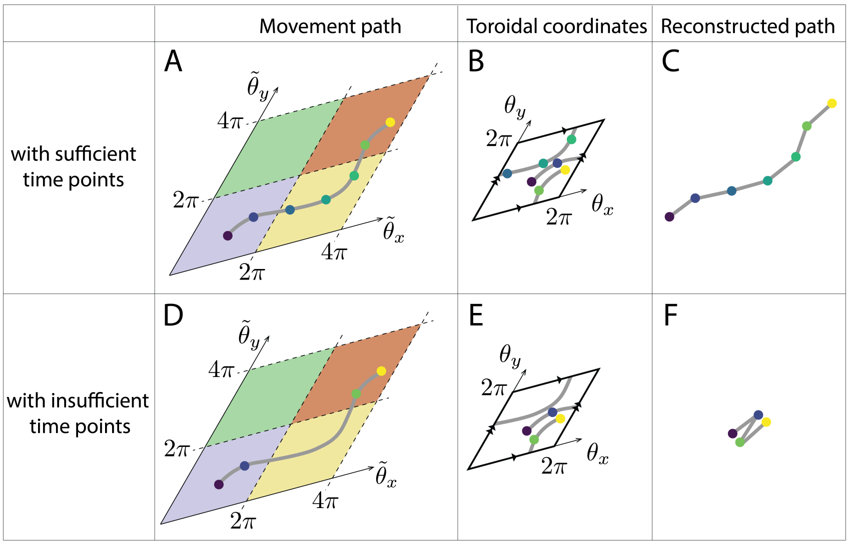


**SI Figure 10.** Having insufficient time points leads to poor path reconstruction **A.** The movement path in $\mathbb{R}^2$ is shown in grey, and the time points at which data is collected are indicated via colored circles. Here, the movement path goes from lower left corner to the upper right corner. **B.** The observed toroidal coordinates are shown in colored dots. **C.** The reconstructed path. **D.** Data is collected at four time points indicated by the colored dots. **E.** The observed toroidal coordinates are shown via colored dots. **F**. In this case, the algorithm fails to lift between the 2nd and 3rd time point, resulting in a reconstructed path that does not resemble the original movement path in panel **D**.

In order to analyze the impact of number of time points in the reconstruction error, we performed the following experiments on the simulated grid cell activity from the 1-hole world.

4.2.1. *Impact of movement speed in simulated movements.* When simulating the mouse trajectory in a bounded environment of size $100 \times 100$, the movement speed is controlled by the parameter $s_{\max}$: at every trajectory step, the distance traveled is sampled uniformly from $[0, s_{\max}]$ spatial units. Larger values of $s_{\max}$ produce faster, more spatially diffuse trajectories, while smaller values produce slower, more locally concentrated paths.

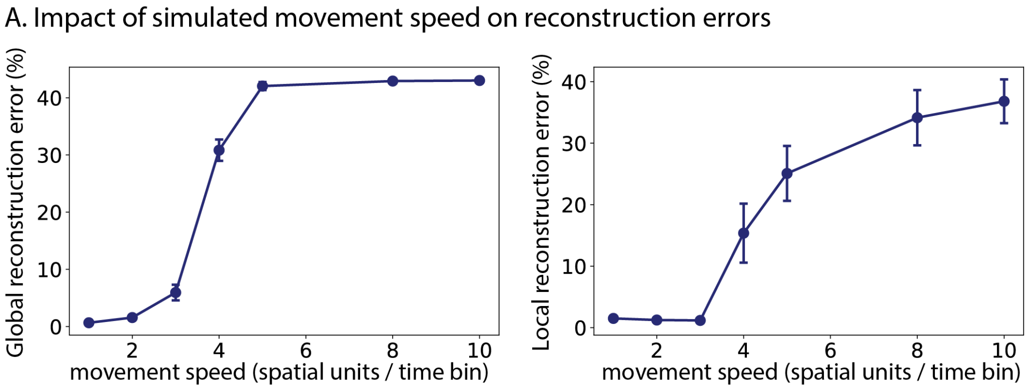


**SI Figure 11.** Global (left) and local (right) reconstruction errors as a function of movement speed $s_{\max}$ (spatial units per trajectory step) in the CAN-simualted data. Each point shows the mean ± std across 5 independent trajectories. Both errors remain near zero for $s_{\max} \leq 3$ and increase sharply for $s_{\max} \geq 4$, indicating that path reconstruction is robust to moderate movement speeds but degrades when the simulated movement speed increases.

To assess the sensitivity of path reconstruction to movement speed, we experimented with $s_{\max} \in \{1, 2, 3, 4, 6, 8, 10\}$ spatial units per trajectory step. For each $s_{\max}$ value, we simulated a movement trajectory of 25,000 steps, simulated grid cell activity (which resulted in 599,999 time bins), and performed path reconstruction using our pipeline. We repeated this process 5 times per $s_{\max}$ value. The global and local reconstruction errors are reported in SI Fig. 11.

While this particular simulation studied movement speed in the simulated trajectory, biologically realistic speeds are unlikely to substantially impair reconstruction performance, since neural processes operate on a much finer timescale. Therefore, this analysis should be considered as a way to examine how the physical distance traveled between sampled time points affects path reconstruction. In real data, the determining factor will be the temporal sampling rate rather than animal speed.

4.2.2. *Impact of uniform temporal subsampling of grid cell activity.* Neural recordings are typically acquired at a fixed sampling rate that does not necessarily coincide with the temporal resolution of the simulations. To assess sensitivity to temporal resolution, a simulated trajectory and grid cell activity, consisting of 599,999 time points, were downsampled to retain every $k$-th time point, for $k \in \{1, 50, 100, 200, 300, 400, 500\}$. We then performed path reconstruction from the downsampled grid cell activity. Each condition was repeated 5 times using independent simulations of a 1-hole environment.

For the local reconstruction errors, we scaled the length of local paths proportionally as $\ell = \lfloor 10{,}000/k \rfloor$, ensuring that each local path corresponds to the same time window regardless of $k$.

Path reconstruction quality was robust across a wide range of downsampling factors (see SI Fig. 12A). Both global and local reconstruction error remained stable for $k \leq 200$. A sharp increase in global reconstruction error occurred at $k = 300$ ($28\% \pm 5\%$). These results indicate that the path reconstruction pipeline tolerates substantial reductions in temporal resolution — up to 200-fold subsampling — before performance degrades, suggesting robustness to practical variations in recording frame rate.

We repeated a similar analysis on the two-dimensional experimental data [4]. Starting from the full-resolution recording of $126,728$ time bins, we uniformly subsampled both the neural activity time bins and ground-truth trajectory by factors of $k \in \{1, 2, 5, 10, 20, 50, 100, 150\}$. To each downsampled dataset, we performed path reconstruction and reported both global and local reconstruction errors. For the local reconstruction errors, recall that the original analysis (without any downsampling) utilized local paths of length 2,000 time bins. The length of the local paths were proportionally shortened to $\lfloor 2,000/k \rfloor$. From SI Fig. 12A (bottom row), one can see that downsampling causes local reconstruction errors to increase beyond $k = 20$.

4.2.3. *Impact of non-uniform temporal subsampling of grid cell activity.* We then tested robustness of the method to irregular temporal sampling. Inter-sample intervals were drawn from a Poisson distribution with mean $\lambda$, and their cumulative sum was used to identify the subsampled time points. We downsampled the grid cell activity with varying levels of $\lambda \in \{1, 50, 100, 200, 300, 400, 500\}$ and performed path reconstruction on the downsampled activity. As in the uniform downsampling analysis, the lengths of local paths were scaled proportionally as $\ell = \lfloor 10{,}000/\lambda \rfloor$.

Similarly to the uniform-downsampling experiments, the global and local reconstruction error remained stable for a wide range of $\lambda$, with performance degrading starting at $\lambda = 200$ (see SI Fig. 12B, top). Compared to uniform downsampling (SI Fig. 12A, top), the degradation onset is somewhat earlier.

We repeated a similar analysis on the two-dimensional experimental data [4]. For each $\lambda \in \{1, 2, 5, 10, 20, 50, 100, 150\}$, we generated 5 independent subsampling of the recorded neural activity by drawing inter-sample intervals from a Poisson distribution with mean $\lambda$. We then performed path reconstruction on each subsampled dataset.

For the analysis of local reconstruction error, we scaled the local path length proportionally as $\ell_{\text{seg}} = \lfloor 2000/\lambda \rfloor$ as before. Similarly to the uniform downsampling analysis, both errors remained stable up to $\lambda = 20$ (see SI Fig. 12B, bottom). Beyond $\lambda = 20$, both global and local reconstruction errors increased gradually. This analysis indicates a modest degradation in local path reconstruction as the neural recording becomes more sparse.

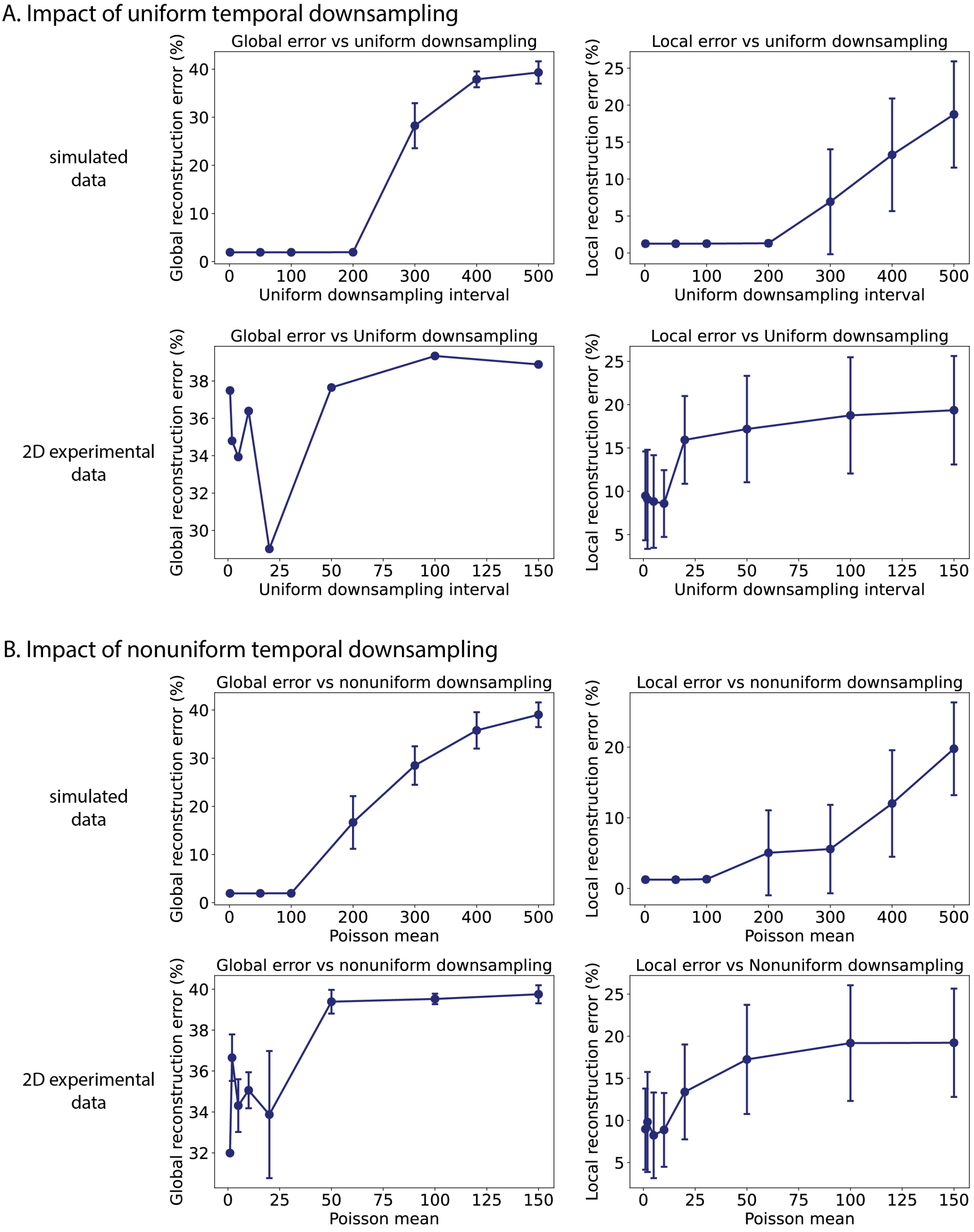


**SI Figure 12.** Impact of temporal subsampling on reconstruction errors. **A.** Global (left) and local (right) reconstruction errors as a function of uniform downsampling interval $k$, for simulated data (top row) and two-dimensional experimental data (bottom row). **B.** Errors as a function of non-uniform (Poisson) subsampling with mean interval $\lambda$. Figures show that simulated data can tolerate up to 200-fold temporal subsampling in the simulated dataset and 20-fold temporal subsampling in the experimental dataset.

4.3. **Error accumulation for long paths and impact of experiment duration.** In the analysis of two-dimensional experimental data [4], the reconstruction of the rat's full trajectory failed to resemble its actual movement, even though shorter segments of the same trajectory were reconstructed faithfully. This discrepancy has two possible sources. First, experimental recordings are noisy, and this noise can cause the path-lifting algorithm to make errors. Second, and more importantly, even a small number of local errors can accumulate, producing a global reconstruction whose shape differs substantially from the original path.

A reconstruction can be a decent reconstruction on local paths and still be globally distorted, because errors during path reconstruction can propagate. SI Fig. 13 illustrates how the two types of lifting error (main text, Fig. 8) distort the global shape of a reconstructed path, even when each local segments is lifted correctly except at the segment boundaries. When the first type of error occurs, toroidal coordinates that should be lifted to the same tile are instead lifted to different tiles. This causes the reconstructed path to be more stretched out than the true movement path. When the second type of error occurs, toroidal coordinates that should be lifted to different tiles are instead placed in the same tile, leading to an overly compressed reconstruction. In both cases, errors occur at only a small number of points, yet the resulting reconstruction differs in global geometry from the true movement path.

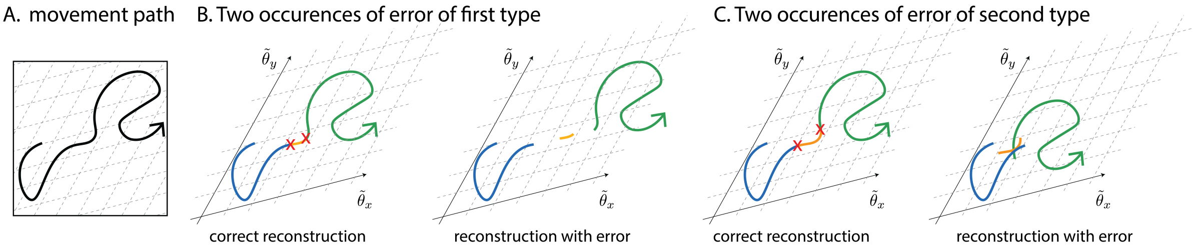


**SI Figure 13.** A small number of lifting errors can distort the global shape of a reconstructed path. Each panel shows an example movement path and the two types of lifting error that can occur during reconstruction. Throughout, the dotted parallelograms indicate the tiles produced by the covering map $p : \mathbb{R}^2 \to S^1 \times S^1$. Each parallelogram is mapped onto a torus under $p$. **A**. An example movement path. **B**. Errors of the first type (over-stretching). Assume that each colored segment of the path is reconstructed correctly, except at the two locations marked by red crosses. At those points, the toroidal coordinates should have been lifted into the same tile as the neighboring segment, but the algorithm lifted them into different tiles. (Left) The correct reconstruction. (Right) The reconstruction produced under this error, which is more stretched out than the correct reconstruction. **C**. Errors of the second type (over-compression). Conversely, assume that at the two marked locations the toroidal coordinates were lifted into the same tile, when they should have been lifted into distinct tiles. (Left) The correct reconstruction. (Right) The reconstruction produced under this error, which is more compact than the correct reconstruction.

We observe this form of error accumulation in the two-dimensional experimental data. In SI Fig. 14A, we highlight a portion of the rat's movement path consisting of three consecutive 20-second segments (labeled 2, 3, 4). SI Fig. 14B shows the reconstruction of the highlighted portion; its geometry is clearly distorted relative to the original. For example, in the original path, segment 3 (yellow) visits regions occupied by segment 1 (navy) and segment 2 (purple), whereas in the reconstruction these segments no longer overlap.

The source of the distortion becomes apparent in SI Fig. 14C, which shows each segment together with its reconstruction. Each segment is reconstructed faithfully locally: the per-segment shape is preserved. However, the reconstruction of segment 3 appears to have been lifted into a different set of tiles than it should have been, producing a long tail (teal highlight in middle and bottom rows) that pulls segment 4 away from segments 2 and 3 in the global reconstruction. A handful of such lifting errors is sufficient to destroy the global geometry.

These observations highlight that even when local reconstruction errors are small, their cumulative effect can lead to large global discrepancies. Therefore, it is important to evaluate both the local reconstruction error, which measures fidelity of reconstruction within short segments,

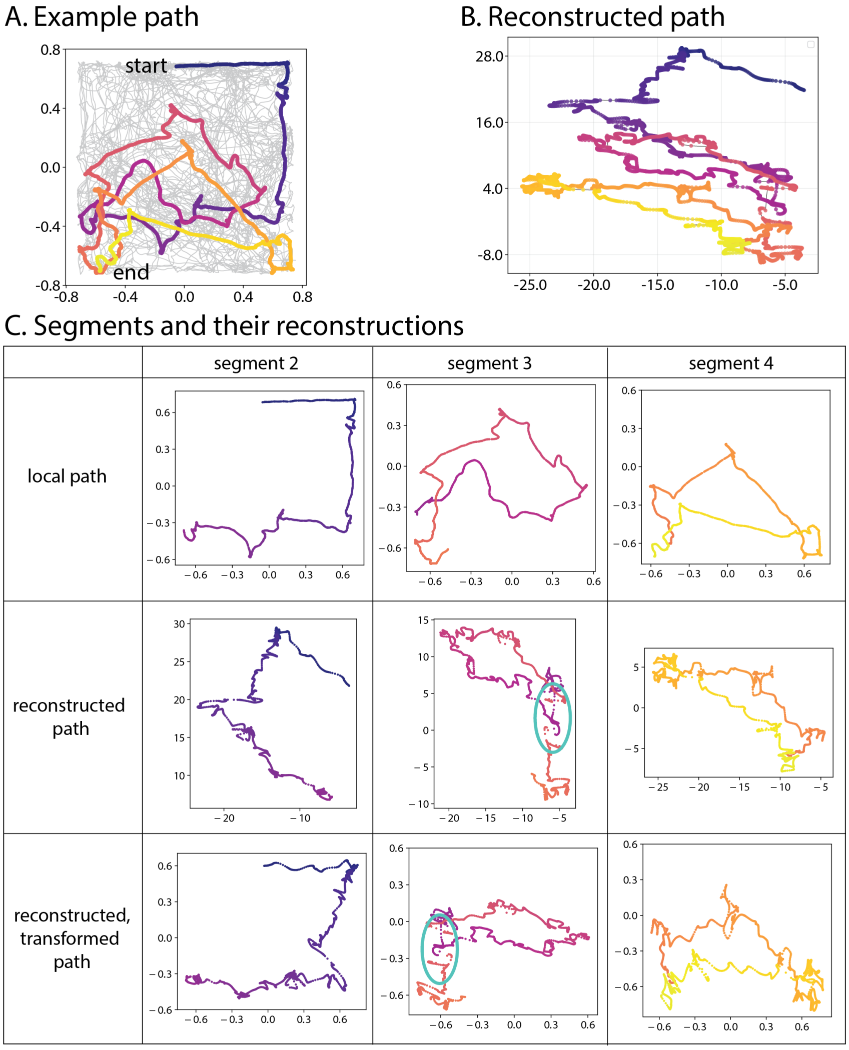


**SI Figure 14.** Error accumulation in the two-dimensional experimental data [4]. We highlight a portion of the rat's trajectory consisting of three consecutive 20-second segments (labeled 2, 3, 4), and compare its reconstruction segment-by-segment and as a whole. The rat moved from navy to purple to yellow across the three segments. **A**. The full movement path in $\mathbb{R}^2$ (gray), with segments 2–4 highlighted. **B**. The reconstruction of the highlighted portion. Its global geometry is clearly distorted relative to the original: for example, segment 4 (yellow) visits regions occupied by segments 2 (navy) and 3 (purple) in the original path, but does not intersect their reconstructions here. **C**. Segment-by-segment comparison. For each of segments 2, 3, and 4: (top row) the original local path, (middle row) its reconstruction, and (bottom row) the reconstruction after aligning to the original coordinate frame. Each segment's local reconstruction faithfully preserves the per-segment shape. The reconstruction of segment 3, however, contains a long "discontinuous" tail (teal highlights in middle and bottom rows), indicating that a portion of the segment may have experienced the first type of error. This lengthened displacement in segment 3 propagates forward, causing segment 4 to be reconstructed in a region that no longer overlaps with segments 2 and 3, thereby distorting the global shape shown in panel B.

and the global reconstruction error, which captures long-range consistency across the full trajectory. A low local error and a high global error indicates that the pipeline is reconstructing each short segment faithfully, but that the errors are accumulating in a way that causes the overall shape of the reconstructed path to differ from the original movement path.

4.3.1. *Impact of duration of experiment.* The length of the experiment can impact the reconstruction quality in several ways. If the recording is too short, the toroidal structure in grid cell population activity may not be sufficiently clear, leading to poor toroidal coordinate computation or failure of computation. However, if the experiment is long enough to yield reliable toroidal coordinates but short enough that the path requires few non-trivial lifts, the reconstruction quality will be high.

In principle, longer experiments should produce clearer toroidal structure and therefore better path reconstructions. However, in the presence of noise, longer experiments are also prone to error accumulation during path lifting. These two effects – improved toroidal coordinates and accumulated lifting errors – compete. To determine which effect dominates, we performed the following experiments on both simulated and experimental data.

Using simulated data on 1-hole environment (full simulation: 599,999 time bins), we truncated the grid cell activity to the first 1,000, 2,000, 3,000, 5,000, 10,000, 20,000, and 30,000 time bins and performed path reconstruction on each truncation. Global and local reconstruction errors were computed over $n = 5$ independent simulations. Local reconstruction errors were evaluated using segments of length 10,000. Both global and local reconstruction errors decreased with longer durations (SI Fig. 15, top row), suggesting that in the simulated setting, the dominant effect is improved toroidal coordinate quality.

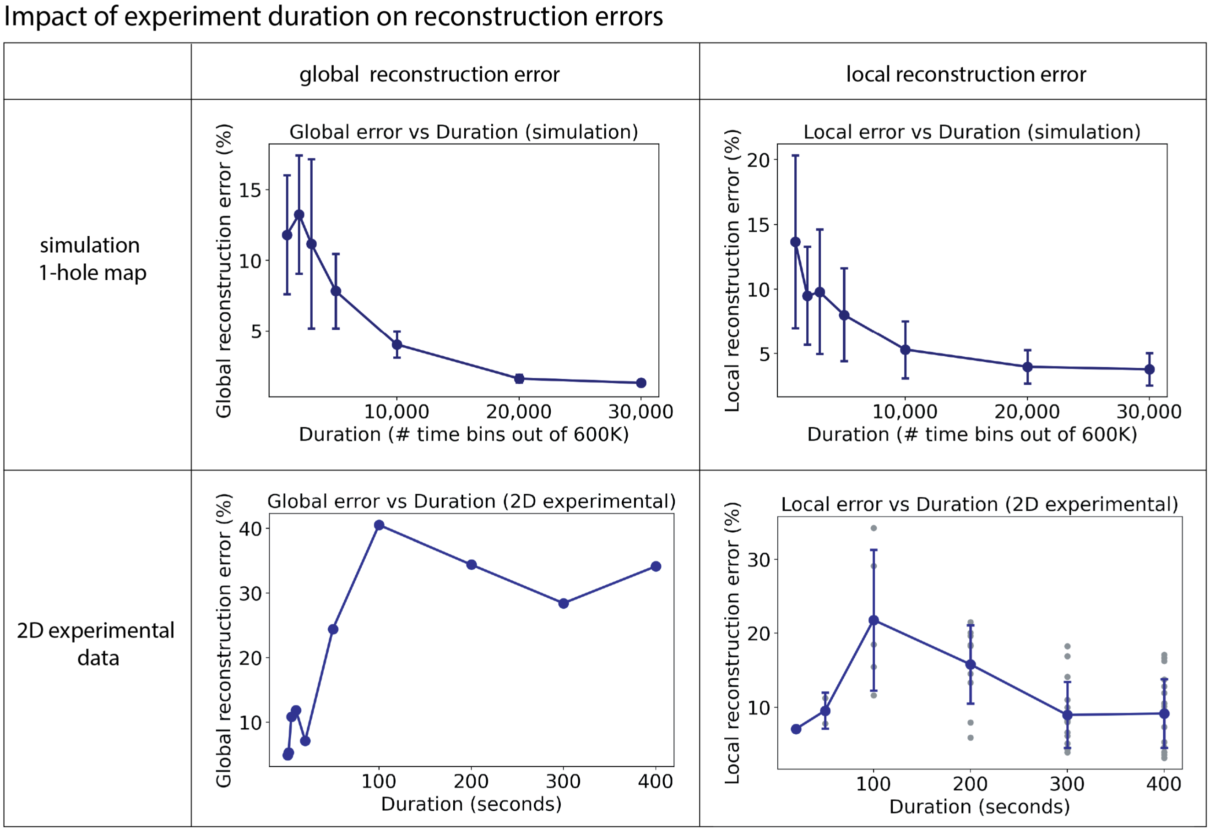


**SI Figure 15.** Impact of experiment duration on reconstruction errors. Global (left) and local (right) reconstruction errors for simulated data (top) and two-dimensional experimental data (bottom).

We repeated the analysis on the two-dimensional experimental data ([4], rat R, module 1, day 2, open-field session; full recording: 21.1 minutes, 126,728 time bins). Grid cell activity was truncated to the first 0.5, 1, 2, 5, 10, 20, 50, 100, 200, 300, and 400 seconds. Because local reconstruction errors were computed using 20-second segments, truncations shorter than 20 seconds were excluded from the local error analysis. The toroidal coordinate computation failed for the 0.5-second truncation.

In contrast to the simulated results, the global and local reconstruction errors exhibited opposite trends (SI Fig. 15, bottom row). The local reconstruction error decreased with duration, consistent with the expectation that longer recordings yield better toroidal coordinates. However, the global reconstruction error increased with duration, likely due to accumulation of lifting errors over longer trajectories.

4.4. **Impact of number of neurons.** The number of simultaneously recorded neurons can affect reconstruction quality. To quantify this effect, we subsampled the simulated grid cell population (2,464 neurons total) by randomly selecting $n \in \{50, 100, 200, 500, 1000, 2000, 2464\}$ neurons and performing the full reconstruction pipeline on the reduced population. Each condition was repeated 5 times with independent random draws of neurons. Note that this differs from simulating grid cell activity using a smaller network: the grid cell activity for 2,464 neurons is simulated using the CAN-model, and the subsampling of neurons is performed on the simulated 2,464 neurons.

We carried out the same analysis on the two-dimensional experimental data of Gardner et al. [4], subsampling $n \in \{10, 20, 40, 60, 80, 100, 111\}$ neurons.

SI Fig. 16 reports the results. In the simulated data, the toroidal coordinates could not be computed for $n = 50$ and $n = 100$, so we report the result only for $n = 200$ and higher. When there were 200 or more neurons, both reconstruction errors were quite low and remained stable. In the experimental data, both global and local reconstruction errors decreased with increased number of neurons.

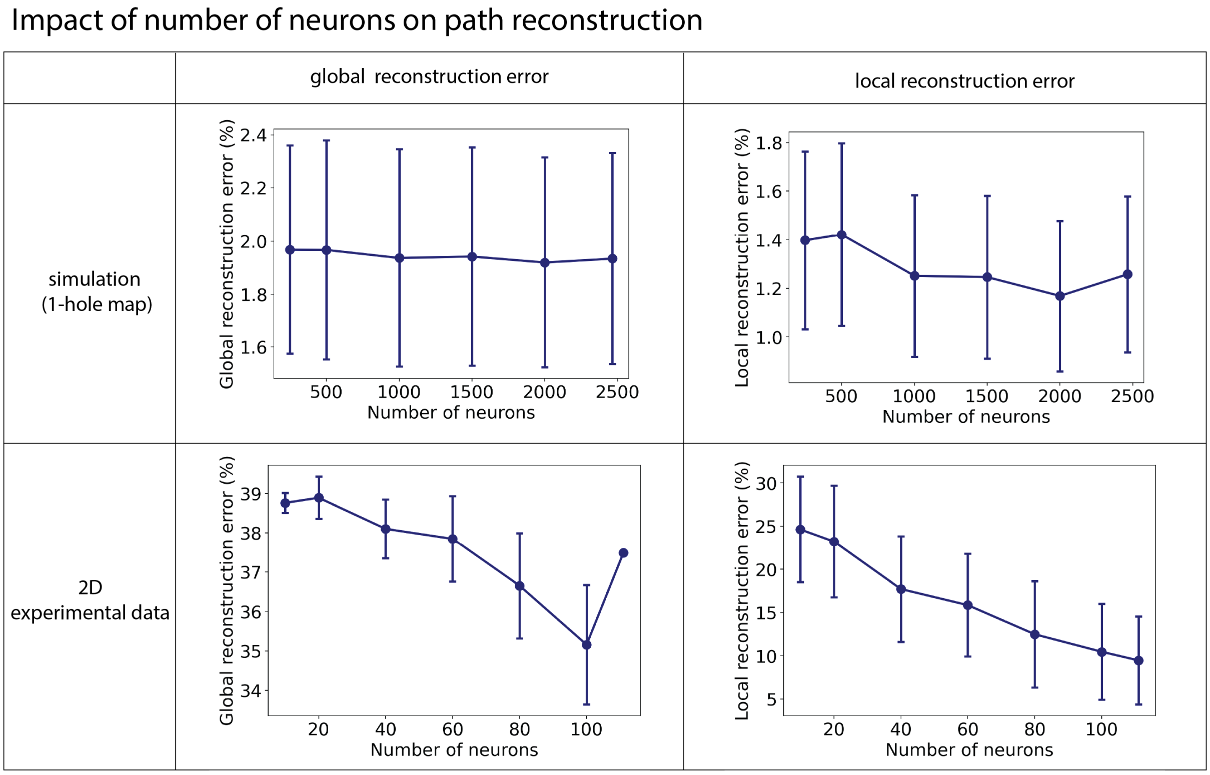


**SI Figure 16.** Impact of neuron count on reconstruction errors. Global (left) and local (right) reconstruction errors as a function of the number of neurons used for path reconstruction, for simulated data (top) and two-dimensional experimental data (bottom).

4.5. **Impact of metric.** The persistent cohomology computation is based on a notion of dissimilarity between population vectors, which can be computed via various choices of metric. Furthermore, the toroidal coordinates can be computed using two distinct methods (DREiMac [7] and cohomological decoding from Gardner et al. [4]). To assess the sensitivity to these choices, we compared the performance of path reconstruction across the two toroidal coordinate computation method and three dissimilarity metrics — cosine, Euclidean, and correlation — on both simulated and experimental data.

For the simulated data, we observed no significant difference between the two toroidal coordinate computation methods and cosine and Euclidean dissimilarity (SI Fig. 17A). Correlation dissimilarity resulted in noticeably higher errors.

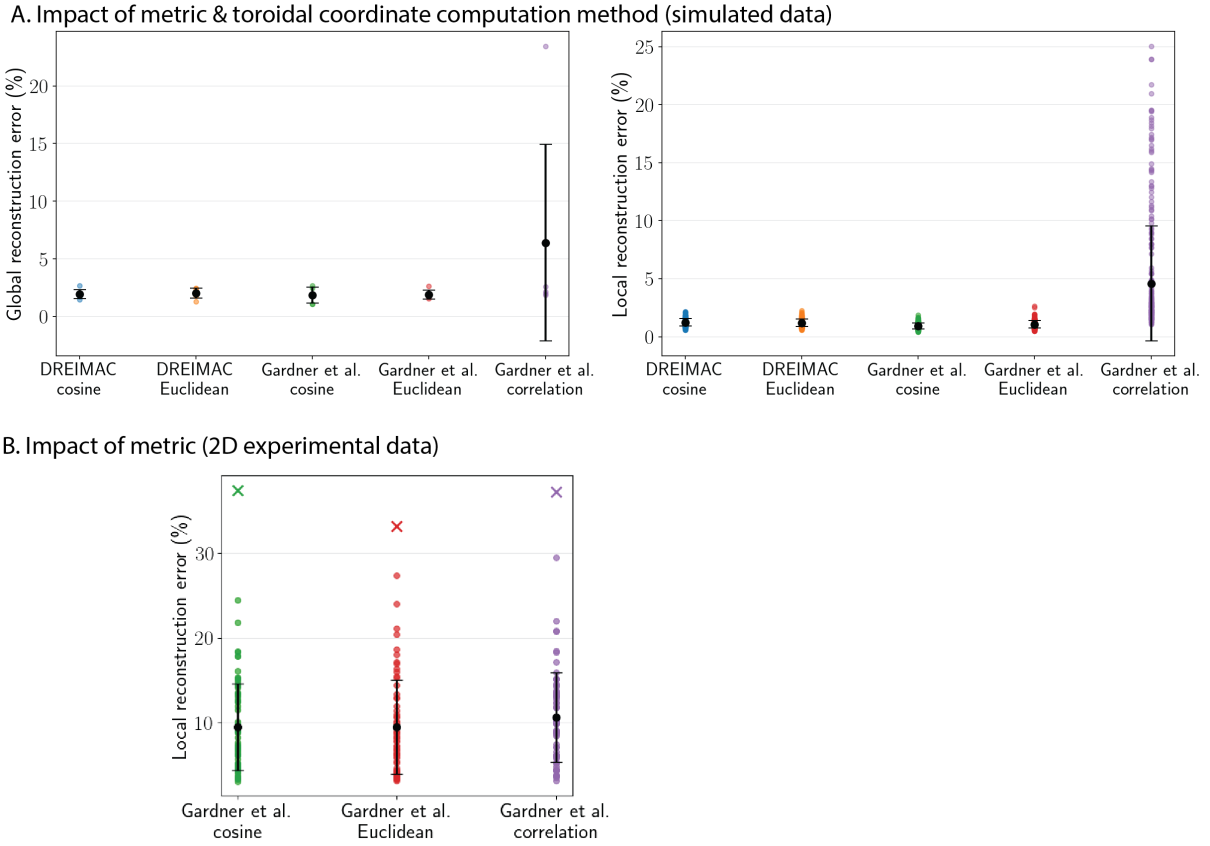


**SI Figure 17.** Impact of toroidal coordinate computation methods (DREiMac [7] vs Gardner et al. [4]) and dissimilarity metric on reconstruction errors. For simulated data, the default method is to use DREiMac with Euclidean dissimilarity. For experimental data, we use the toroidal coordinate computation from Gardner et al. [4] with cosine dissimilarity. **A.** On the CAN-simulated data on one-hole environment, we compared the performance of path reconstruction using two methods of computing toroidal coordinates (DREiMac and Gardner et al. [4]) and three dissimilarity metrics (cosine, Euclidean, and correlation) over five independent simulations. Both global (left) and local (right) reconstruction errors were similar, except for using correlation dissimilarity. **B.** Global ($\times$) and local ($\circ$) reconstruction errors for two-dimensional experimental data [4].

For the two-dimensional experimental data (Gardner et al. [4], rat R, module 1, day 2, open-field), where we utilized the cohomological decoding of Gardner et al. [4], all three dissimilarity metrics yielded comparable global and local reconstruction errors (SI Fig. 17B).

Together, these results suggest that the pipeline is relatively robust to the choice of dissimilarity metric and the toroidal coordinates computation method.

4.6. **Impact of noise in toroidal coordinates.** The path-lifting algorithm takes as input a sequence of toroidal coordinates $\{\Theta(t)\} = \{(\theta_x^t, \theta_y^t)\}$ that have been computed via persistent cohomology. To characterize how sensitive the reconstruction is to noise in toroidal coordinates, we perturbed the toroidal coordinates with additive Gaussian noise and measured the resulting reconstruction quality.

Specifically, we first simulated the grid population activity in the one-hole environment and computed the toroidal coordinates. Given toroidal coordinates $(\theta_x^t, \theta_y^t)$ at time $t$, we replaced the coordinates by $(\theta_x^t + n_x,\ \theta_y^t + n_y)$, where $n_x$ and $n_y$ were independently drawn from $\mathcal{N}(0, \sigma^2)$. The perturbed values were then wrapped to $[0, 2\pi)$. We considered $\sigma \in \{0, 0.1, 0.2, 0.3, 0.5, 0.75, 1.0, 1.5, 2.0, 3.0\}$ and repeated each condition 5 times.

SI Fig. 18A visualizes the perturbed toroidal coordinates for $\sigma \in \{0, 0.5, 1, 2\}$. As $\sigma$ increases, the coordinate structure becomes progressively less well-defined. SI Fig. 18B reports the global and local reconstruction errors as a function of $\sigma$. Both error measures remained low for $\sigma \leq 0.3$

and increased sharply around $\sigma = 0.5$. At $\sigma \geq 1.0$, errors plateaued at high values, indicating that the toroidal coordinate signal had been largely destroyed.

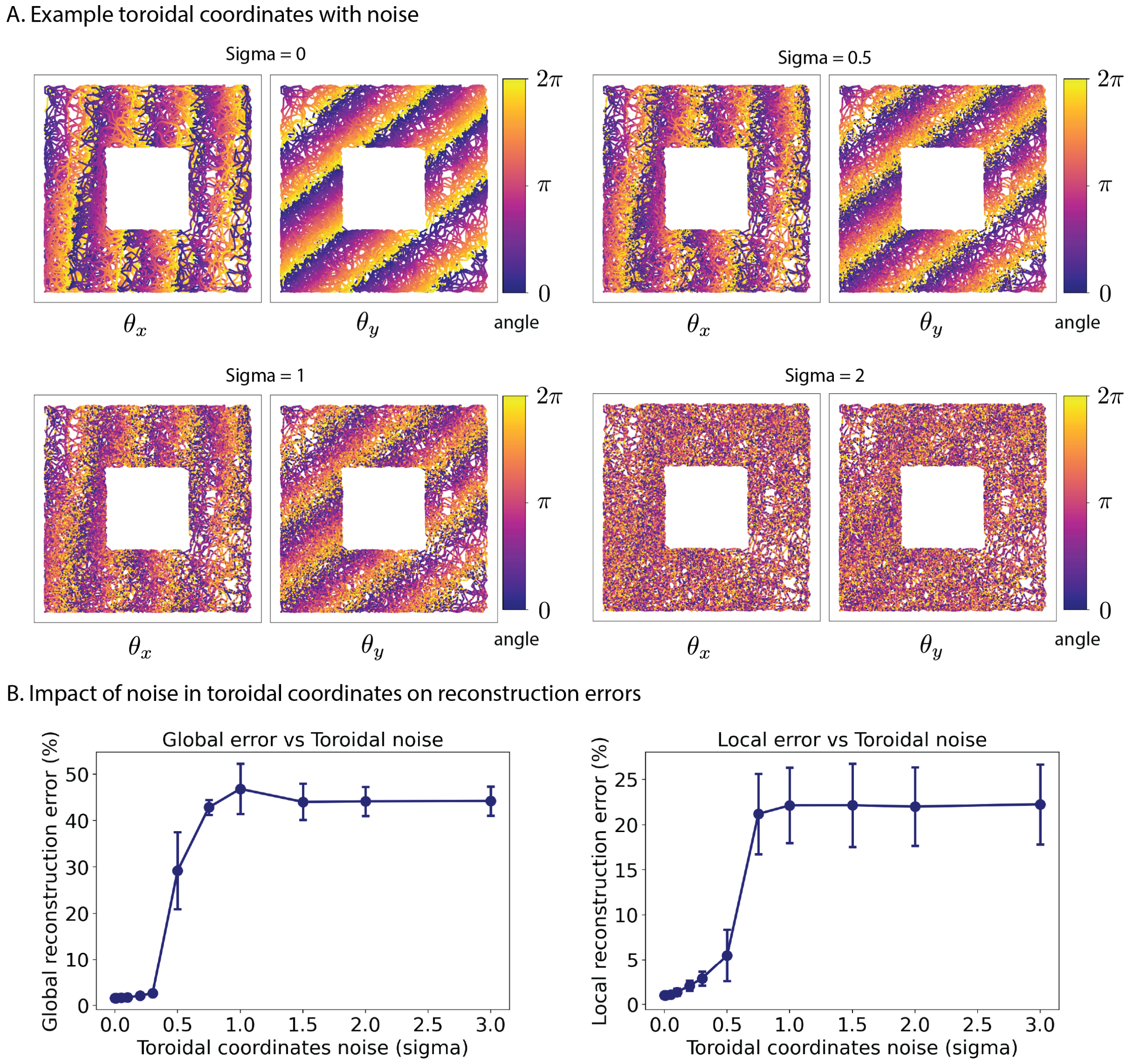


**SI Figure 18.** Impact of noise in toroidal coordinates on reconstruction error in simulated data (1-hole environment, $n = 5$ repeats). **A.** Example toroidal coordinate visualizations at noise levels $\sigma \in \{0, 0.5, 1, 2\}$. **B.** Global (left) and local (right) reconstruction errors as a function of $\sigma$.

4.7. **Impact of smoothing reconstructed paths from experimental data.** Paths reconstructed from two-dimensional experimental data (rat R, module 1, day 2, OF; [4]) contain a lot of jitter, as illustrated in Fig. 7 of main text, as well as SI Fig. 19A. One may thus choose to report the reconstructed path after smoothing the path to a certain degree. For example, one might choose to apply Gaussian smoothing with varying standard deviations ($\sigma$ = 0, 1, 5, 10, 20, 30, 50). Applying such smoothing preserves the overall trajectory shape while reducing jitter. Such smoothing has minimal impact on both global and local reconstruction errors (see SI Fig. 19B, C).

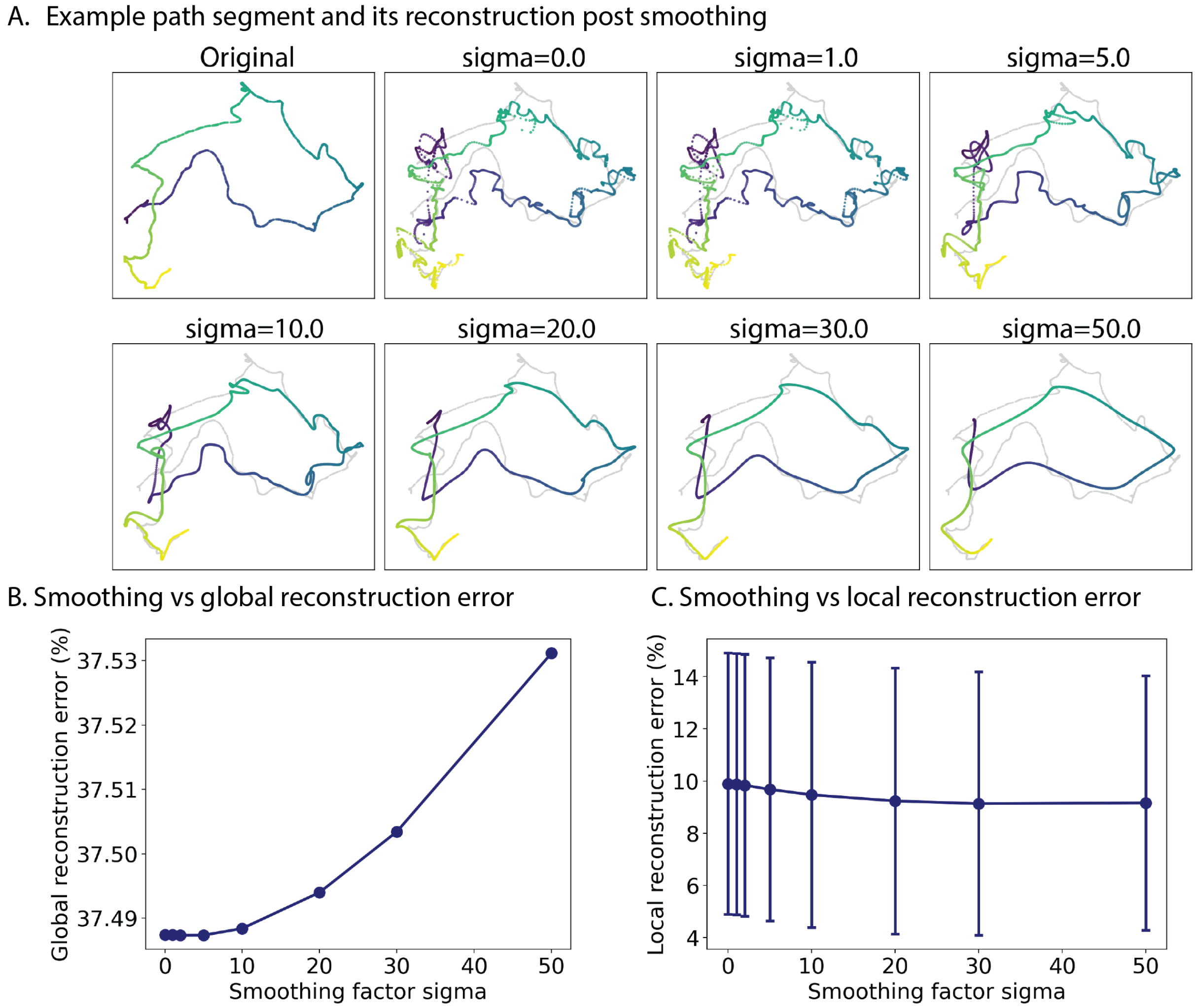


**SI Figure 19.** Impact of smoothing on an example reconstructed path from two-dimensional experimental data (rat R, module 1, day 2, OF; [4]). **A.** An example local path segment (left, "Original") and its reconstruction after applying Gaussian smoothing with varying standard deviations ($\sigma$ = 0, 1, 5, 10, 20, 30, 50). **B.** Moderate levels of smoothing (up to $\sigma = 50$) has minimal impact on global reconstruction error, which stays around 37%. **C.** Increased smoothing improves local reconstruction error slightly.

372

### 5. Supplementary Figures

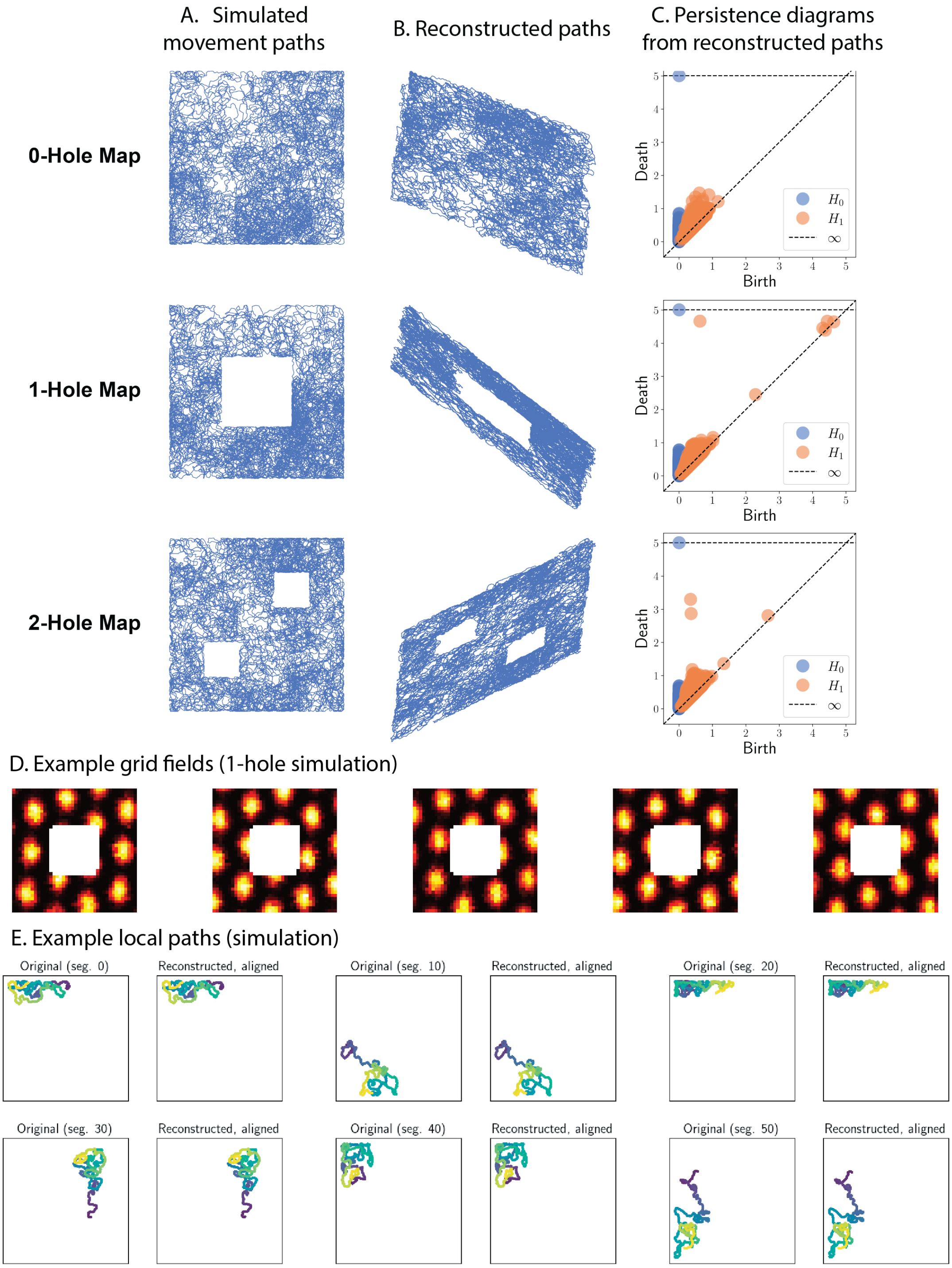


**SI Figure 20.** Path lifting on simulated grid cell activity preserves the global topology of the environment. **A**. Simulated movement trajectories in environments with 0, 1, and 2 holes. **B**. Reconstructed paths from simulated grid cell activity. **C**. To show that the number of holes in the environment can be recovered from the reconstructed path, we computed pairwise Euclidean dissimilarity between every pair of points in the reconstructed path and computed persistent homology of the Vietoris-Rips filtration. The persistence diagrams computed on the reconstructed path recover the correct number of holes in the environment. **D.** Example grid fields from simulated grid cells. **E.** Example local paths corresponding to $10,000$ time bins and their reconstructions, post alignment. The original trajectory corresponds to $599,999$ time bins.

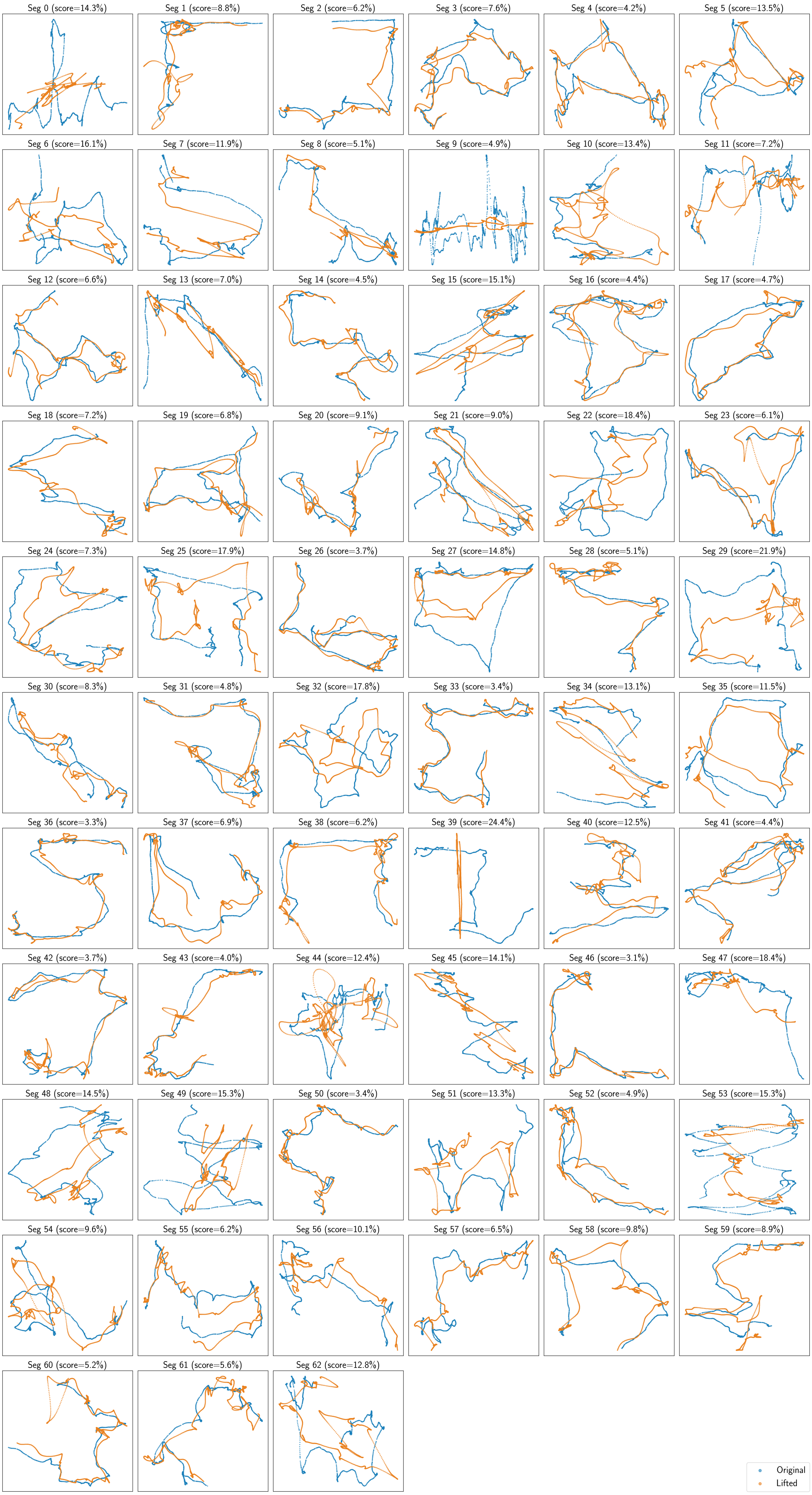


**SI Figure 21.** Complete local path reconstruction for two-dimensional experimental data from [4]. In

### References

[1] Gunnar Carlsson. "Topology and Data". In: *Bulletin of The American Mathematical Society - BULL AMER MATH SOC* 46 (Apr. 2009), pp. 255–308. DOI: 10.1090/S0273-0979-09-01249-X.

[2] Herbert Edelsbrunner, John Harer, et al. "Persistent homology-a survey". In: *Contemporary mathematics* 453.26 (2008), pp. 257–282.

[3] Edelsbrunner, Letscher, and Zomorodian. "Topological persistence and simplification". In: *Discrete & computational geometry* 28 (2002), pp. 511–533.

[4] Richard J. Gardner, Erik Hermansen, Marius Pachitariu, Yoram Burak, Nils A. Baas, Benjamin A. Dunn, May-Britt Moser, and Edvard I. Moser. "Toroidal topology of population activity in grid cells". In: *Nature* 602.7895 (Feb. 2022). Publisher: Nature Publishing Group, pp. 123–128. ISSN: 1476-4687. DOI: 10.1038/s41586-021-04268-7.

[5] Robert Ghrist. "Barcodes: the persistent topology of data". In: *Bulletin of the American Mathematical Society* 45.1 (2008), pp. 61–75.

[6] J. Munkres. *Topology*. Pearson Modern Classics for Advanced Mathematics Series. Pearson, 2017. ISBN: 9780134689517.

[7] Jose A. Perea, Luis Scoccola, and Christopher J. Tralie. "DREiMac: Dimensionality Reduction with Eilenberg-MacLane Coordinates". In: *Journal of Open Source Software* 8.91 (2023), p. 5791. DOI: 10.21105/joss.05791.
